# Genomic insights into the adaptive and convergent evolution of *Leuciscus waleckii* inhabiting extremely alkaline environments

**DOI:** 10.1101/2022.05.15.492028

**Authors:** Zhixiong Zhou, Junyi Yang, Hongzao Lv, Tao Zhou, Ji Zhao, Huaqiang Bai, Fei Pu, Peng Xu

## Abstract

*Leuciscus waleckii* is widely distributed in Northeast Asia and has high economic value. Different from its freshwater counterparts, the population in Lake Dali Nur has a strong alkalinity tolerance and can adapt to extremely alkaline–saline water with bicarbonate over 50 mmol/L (pH 9.6), thus providing an exceptional model with which to explore the mechanisms of adaptive evolution under extreme alkaline environments. Here, we assembled a high quilty chromosome-level reference genome for *L. waleckii* from Lake Dali Nur, which provides an important genomic resource for the exploitation of alkaline water fishery resources and adaptive evolution research across teleost fish. Notably, we identified significantly expanded long terminal repeats (LTRs) and long interspersed nuclear elements (LINEs) in *L. waleckii* compared to other Cypriniformes fish, suggesting their more recent insertion into the *L. waleckii* genome. We also identified expansions in genes encoding gamma-glutamyltransferase, which possibly underlie the adaptation to extreme environmental stress. Based on the resequencing of 85 *L*.*waleckii* individuals from divergent populations, the historical population size of *L*.*waleckii* in Lake Dali Nur dramatically expanded in a thousand years approximately 13,000 years ago, and experienced a cliff recession in the process of adapting to the alkaline environment of Lake Dali Nur approximately 6,000 years ago. Genome scans further revealed the significant selective sweep regions from Lake Dali Nur, which harbour a set of candidate genes involved in hypoxia tolerance, ion transport, acid-base regulation and nitrogen metabolism. In particular, 5 alkali population specific nonsynonymous mutations were identified in CA15 gene copies. In addition, two sites with convergent amino acid mutation were detected in the RHCG-a gene among several alkali environment adapted Cypriniformes fish, this mutation may increase the NH_3_ excretion rate of the RHCG channel. Our findings provide comprehensive insight into the genomic mechanisms of *L. waleckii* and reveal their adaptative evolution under extreme alkaline environments.

## Introduction

Diverse extreme environments, including deserts, plateaus, deep seas and saline-alkali lakes are important components of the global ecosystem. In the process of survival under these extreme environments, species generally undergo unique adaptive evolution, which popular research topic in the field of biological evolution^1^. Many studies have shown that the selection pressure caused by extreme environments may change the protein coding, expression pattern, copy number or molecular function of several core genes in adaptive species^2-5^. Moreover, different species that adapt to the same extreme environment may adopt convergent genetic evolution strategies^6^. These cases of classic adaptive evolution could help us understand the mystery of biological evolution and provide new avenues for reasonably developing genetic resources for rare extreme environmental adaptive species.

Alkalization and salinization, which are critical threats to inland lakes and freshwater fishery resources, are currently widespread and occur at an unprecedented rate under intensifying global warming^7^. Usually, the pH of these alkaline-saline lakes is higher than 9.0, and the salinities may approach approximately 50% for seawater^8^. This extremely alkaline environment could disrupt the acid-base balance, inhibit the excretion of nitrogenous waste and disturb the osmotic pressure regulation of alkaline-adapted fish. Although the effects of these extreme environmental factors on the reproduction and growth of freshwater fish are lethal, several fish can naturally survive in alkaline-saline lakes. Therefore, the adaptive evolutionary mechanism of these recurring alkaline-adaptive fish has long been of interest to evolutionary biologists^9-11^. Generally, to avoid elevated blood pH due to respiratory alkalosis in alkaline-saline lakes, teleosts regulate the blood pH through reversible CO_2_ hydration/dehydration reactions catalysed by carbonic anhydrase (CAs)^12^. Additionally, several important ion transport channels are also involved in the regulation of acid-base balance, such as Cl^-^/HCO_3_^-^ and Na^+^/H^+^ exchangers across the gill, which has been confirmed in Lahontan Cutthroat trout (*Oncorhynchus clarki henshawi*), rainbow trout (*Oncorhynchus mykiss*) and naked carp (*Gymnocypris przewalskii*)^13-15^. To deal with the inhibition of ammonia excretion under highly alkaline environment, freshwater teleosts have a variety of coping strategies, such as reducing the metabolic rate, actively excreting ammonia in the gill, converting accumulated nitrogenous waste to nontoxic glutamine or free amino acids and synthesizing urea^16-19^. The classic case is Magadi tilapia (*Alcolapia grahami*), which inhabits Lake Magadi, with high pH (∼10) and salinity (∼60 % of seawater)^18^. Transcriptome evidence showed that Magadi tilapia had a functional ornithine-urea cycle pathway in the gills, which was conducive to increasing nitrogenous waste efficiency by excreting urea^20^. Therefore, exploring the adaptive evolution of fish that can survive in an extremely alkaline environment can help to exploit the fishery potential in alkaline-saline lakes and provide new perspectives on the genetic mechanism of important physiological regulation in teleost fish^21^.

Amur Ide (*Leuciscus waleckii*) is a common freshwater fish in Northeast China with high economic value and is a food source for migrating birds from Siberia ^22^. As an extreme example, a special Amur Ide population can survive in the extreme alkaline environment of Lake Dali Nur located on the eastern Inner Mongolia Plateau, which is a typical saline-alkaline lake with high concentrations of carbonate salts that is lethal to most freshwater teleosts (Supplymentary File1: Fig S1). Currently, the pH value of Lake Dali Nur ranges from 8.25 to 9.6, with an alkaline content (ALK) over 50 mg/L and a salinity of approximately 6 ‰^23^. Based on geological and biological evidence, the prevailing view is that the Amur Ide population in Lake Dali Nur was a freshwater fish that evolved quickly in the past several thousand years and has developed great tolerance to high alkalinity^24^. It is a model species used to explore the adaptation of teleosts to extreme alkaline environments because it has different populations living in alkaline and freshwater areas^11^. Hence, scientists have been interested in the mechanism of its microevolution in the past decade, as the species rapidly evolved to survive rapid paleoenvironmental changes since the early Holocene^10,24-27^. However, the genomic signature and key adaptive evolutionary loci underlying the tolerance to high alkali conditions should be further explored, particularly through comparative genetic analysis between alkaline and freshwater Amur Ide populations.

Here, we present a high-quality chromosome-level genome of *L. waleckii* inhabiting the extremely alkaline waters of Lake Dali Nur. Comparative genomics analysis revealed a series of adaptive evolution events in the *L. waleckii* genome in response to an extremely alkaline environment. Microevolution scanning of alkaline and freshwater populations explained the physiological regulatory mechanism and revealed several key adaptive evolutionary loci in the Lake Dali Nur Amur Ide population.

## Results and discussion

### Genome assembly and annotation

A high-quality chromosome-level genome is needed for the downstream analysis of adaptive microevolution^28^. Using the established method, the genome size was evaluated to be approximately 1,125.03 Mb, the heterozygous rate and repeat rate were evaluated as 0.56% and 57.61% respectively by 17-mer analysis (Supplymentary File1: Fig S2 and Table S3) ^29^. Using the PacBio platform, we sequenced the genomes of Amur Ide in Lake Dali Nur (See Supplementary File 1). The assembled genome spanned 1,103 Mb, with a contig N50 length of 1.52 Mb (Supplymentary File1: Tables S4). Genome annotation showed that the Amur Ide genome comprises approximately 49.92% repetitive sequences (Supplymentary File1: Tables S5 and Supplymentary File2: Table S6), which was comparable to the repeat content of other Cypriniform species genomes^30-32^. In the Amur Ide assembly, we predicted 27,633 protein-coding genes, of which 96.3% of the protein sequences showed similarity to protein sequences in public databases (Supplymentary File2: Table S7, Table S8, Supplymentary File1: Fig S3 and Fig S4). The contigs were then anchored and oriented into a chromosomal-scale assembly using the Hi-C scaffolding approach. Ultimately, we obtained a draft genome assembly of 1,105 Mb in length, with a scaffold N50 value of 39.64 Mb (Supplymentary File1: Table S9). The genome assembly contained 25 chromosomes, with chromosome lengths ranging from 28.42 to 71.37 Mb, and covered 1,020 Mb (92.32%) of the *L*.*waleckii* assembly (Supplymentary File1: Table S9, Fig S5 and Fig S6A). BUSCO analysis showed that the assembly retrieved 96.4% of the conserved single-copy orthologous genes (Supplymentary File1: Table S10). In addition, we mapped Illumina short reads to the L. waleckii reference genome with a mapping ratio of 99.05% and generated 2,707,134 SNPs (Supplymentary File1: Table S11, Table S12). This evidence supported the high-quality assembly of the L. waleckii genome.

### Comparative genomics revealing convergent evolution among alkaline adaptive teleosts

The chromosome synteny comparisons among *L. waleckii, A. nigrocauda* and *C. idella* showed that chromosomes 10 and 22 of *L. waleckii* fused into one chromosome in *A. nigrocauda* and *C. idella* (Fig. 1A and Supplymentary File1: Fig S6B). This chromosome fusion event was also detected in four East Asian cyprinids (silver carp, bighead carp, grass carp, and blunt snout bream)^33^. Our results suggested that this East Asian cyprinid-specific chromosome fusion event occurred later than the differentiation of *L. waleckii* and grass carp (36.2 mya-54.0 mya), which may have led to the evolution and expansion of the East Asian cyprinid clade^33,34^.

**Fig. 1.**
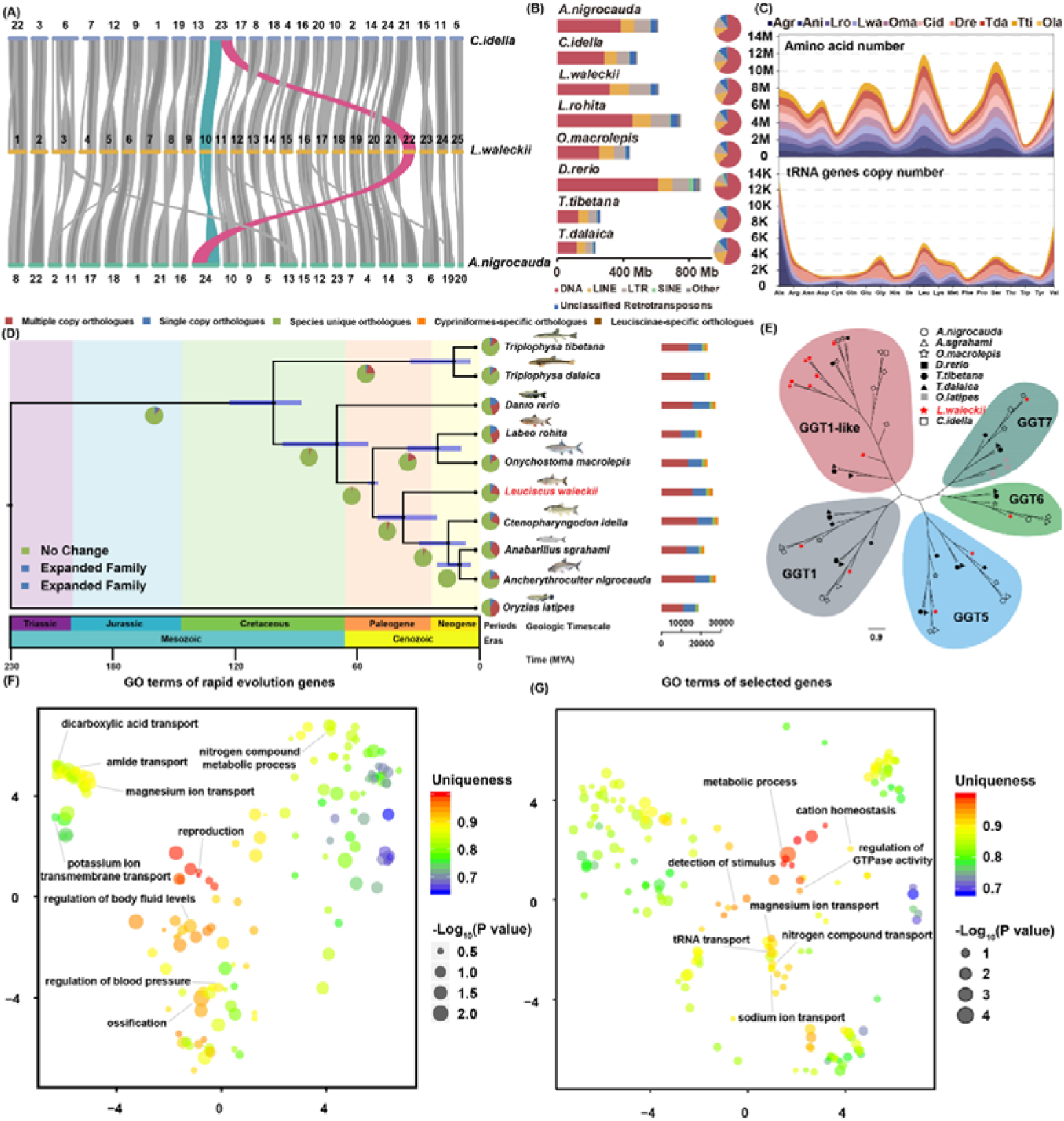
Comparison of evolutionary features of *L. waleckii* and other Cyprinidae species. (A) Genomic collinearity of *L. waleckii* and its related species; (B) Comparison of various transposable elements between *L. waleckii* and related species; (C) Comparison of the number of specific amino acids used in protein-coding and tRNA gene copies between *L. waleckii* and related species; and (D) Divergence times and distribution of different types of orthologues in representative species. Estimated divergence times of representative species based on phylogenomic analysis. The blue bars in the ancestral nodes indicate the 95% confidence intervals of the estimated divergence time (MYA, million years). Different background colours represent the corresponding geological ages. (E) Phylogenetic tree of GGT proteins in vertebrates showing gene expansion in the *L. waleckii* genome; (F) GO terms of positively selected genes summarized and visualized as a REVIGO scatter plot; and (G) GO terms of rapidly evolving genes summarized and visualized as a REVIGO scatter plot. Each circle represents a cluster of related GO terms, with a single term chosen by REVIGO as the cluster representative. Clusters are plotted according to semantic similarities to other GO terms (adjoining circles are most closely related). “Uniqueness” (the negative of average similarity of a term to all other terms) measures the degree to which the term is an outlier when compared semantically to the whole list.

The adaptive evolution of transposable elements (TEs) and gene regulation of the insertion region of TEs may have important evolutionary effects in the process of species adapting to extreme environments^35^. Hence, we used a common protocol to identify TE in *L. waleckii* and 7 Cypriniformes species to compare the TE contents, types and divergence (see Methods). In the *L. waleckii* genome, approximately 598 Mb (54.17% of genome size) were composed of TEs (Supplymentary File2: Table S13). Among the interspersed repeats, the most abundant transposable elements were DNA transposons (28.59% in genome). Retrotransposons were the second most abundant repeat elements (22.85% in genome), including three major families of long terminal repeats (LTRs), long interspersed elements (LINEs), and short interspersed elements (SINEs). The TE contents and their proportions in *L. waleckii* were similarto those in other Cyprinidae fish genomes except for the LTRs and LINEs, in which LTRs were significantly expanded to 11.47% in *L. waleckii* and LINEs were significantly expanded to 10.87% in *L. waleckii* (Figure 1C and Supplymentary File2: Table S13). The insertion time of TEs can be estimated based on their Kimura substitution level. Compared with those in *A. nigrocauda* and *C. idella*, we found that the LTRs and LINEs in *L. waleckii* significantly expanded with 15% divergence rates (Supplymentary File1: Fig S9). In another extremely alkaline environment adapted Cyprinidae fish, *T. dalaica*, the LTRs and LINEs also expanded with 13% divergence rates compared to those of *T. tibetana*, which inhabits a freshwater environment. Hence, the expansion of LTRs and LINEs in two Lake Dali Nur-specific fish may play an important role in their adaptation to an extremely alkaline environment. In addition, we found that the frequency of leucine (Leu) in *L. waleckii* (2.42%) was higher than that in other Cyprinidae species, and the frequency of Leu in *T. dalaica* (2.45%) was also higher than that in *T. tibetana* (2.43%) (Fig. 1D and Supplymentary File2: Table S14). Similarly, by detecting the tRNA gene copy numbers of 10 teleost species, we found that the tRNA gene copy numbers of glutamic acid (Glu) and valine (Val) in *L. waleckii* (569 and 1,196) were larger than those in other Leuciscinae-Culterinae species (38-451 and 51-793), and the tRNA gene copy numbers of Glu and Val in *T. dalaica* (261 and 3,043) were also larger than those in *T. tibetana* (214 and 1,619) (Fig. 1D and Supplymentary File2: Table S15). Therefore, we suggest that two Lake Dali Nur-specific alkaline-adapted fish may use more Leu in the protein-encoding process and have increased the tRNA gene copy numbers of Glu and Val in an extremely alkaline environment.

### Expanded gene families underlying alkaline adaptation of *L. waleckii*

To determine the phylogenetic relationship of *L. waleckii* in Cypriniformes species, we compared ten teleost genomes. Based on 1,635 single-copy orthologues shared among ten fishes, the phylogenetic tree revealed that *L. waleckii* and *C. idella* separated approximately 36.2 mya (confidence interval 19.3 to 49.5 mya), which corroborates the results of other recent studies based on smaller datasets (Fig. 1D and Supplymentary File2: Table S16)^30,31^. Moreover, we uncovered 1,751 *L. waleckii* gene families with expansion and 5,202 families with contraction (Fig. 1D and Supplymentary File1: Table S17). GO enrichment analysis showed that the expanded gene families were mainly involved in cell death, lipid transport, glutathione catabolic processes and chromatin assembly, and the contracted gene families were mainly involved in ion transport, germ cell development, calcium ion homeostasis and cell recognition (Supplymentary File1: Table S17, Fig S8, Fig S9 and Supplymentary File2: Table S18,).

As an important regulator in the glutamine and glutathione metabolic pathway, gamma-glutamyltransferase (GGT) cleaves the gamma-glutamyl bond, releases free glutamate and the dipeptide cysteinyl-glycine, and transfers the gamma-glutamyl moiety to an acceptor amino acid to form a new gamma-glutamyl compound ^36^. In addition, increased plasma GGT has been confirmed to accelerate the rate of ammonia synthesis in blood and plasma ^37^. There are usually 6 to 10 copies of GGT genes in Cyprinidae species, including GGT1, GGT1-like, GGT5, GGT6 and GGT7. However, we identified 14 copies of GGT genes in *L. waleckii*, which were considerably expanded compared with those in other Cyprinidae species (Fig. 1E and Supplymentary File2: Table S20). Of the 14 GGT genes, GGT1-like genes were expanded to 8 copies. The physiological role of GGT in glutamine and glutathione metabolism has been showed in various vertebrates ^38-40^. Under an extreme alkaline environment, converting the accumulated endogenous ammonia into less toxic glutamine, glutathione, free amino acids or urea is an important response mechanism for teleosts to excrete excess ammonia nitrogen waste^17^. Therefore, the expansion of GGT1-like genes in the *L. waleckii* genome might be among the adaptive changes that enhance the synthetic capacity of less toxic glutamine and glutathione.

### Positively selected and rapidly evolving genes in *L. waleckii*

In the process of adapting to extreme environments, natural selection will form selection marks on several important genes, among which the most important types are positive selection genes (PSGs) and rapid evolution genes (REGs)^4^. We identified a set of 131 REGs in the *L. waleckii* lineage, including glutathione peroxidase 7 (*gpx7*), AMP deaminase 2 (*ampd2*), and others (Supplymentary File2: Table S21 and Table S22). In addition, we identified 369 PSGs in *L. waleckii*, including uromodulin-like 1 (*umodl1*), ammonium transporter Rh type A (*rhag*), glutamate receptor ionotropic, delta-1 (*grid1*), and others (Supplymentary File2: Table S23 and Table 24). As an alternative to tallying GO terms, we also examined the nature of GO patterns in the complete set of these REGs and PSGs. By applying the GO clustering tool REVIGO^41^ to the terms associated with the REGs from *L. waleckii*, we found that terms related to reproduction, ossification, and blood circulation had low average similarity (i.e., higher unique; uniqueness > 0.9) (Figure 1F and Supplymentary File2: S25). In addition, several terms related to ion transport, energy metabolism and ammonia nitrogen metabolism had medium average similarity (medium unique; 0.7 < uniqueness < 0.9), which suggests that they were important in the adaptive evolution of extreme alkaline environments^10^. Conversely, in PSGs, terms related to ion transport, metabolic process and ammonia nitrogen metabolism had low average similarity (i.e., higher unique; uniqueness > 0.9), which indicates that a higher proportion of REGs in the *L. waleckii* genome may be involved in the adaptive evolution to an extreme alkaline environment (Figure 1G and Supplymentary File2: Table S26).

### Population genomic analysis of *L. waleckii*

*L. waleckii* was widely distributed in the aquatic ecosystem of Northeast China. Due to the extremely alkaline environment of Lake Dali Nur, the *L. waleckii* population is a marvellous example of rapid adaptive evolution. Eighty-five individuals were resequenced to investigate their genetic variation and the selective signatures of adaptive evolution (Figure S1 and Table S27). After applying stringent quality control criteria, we identified a total of 6,206,224 SNPs and 845,427 insertions/deletions (INDELs) (Supplymentary File1: Table S28). Based on a phylogenetic tree, 85 individuals were grouped into three distinct clades, including the WS-HL group, DL group and YD group (Fig. 2A). Similar results were obtained by PCA and admixture analysis, with the DL and WS-HL populations being completely differentiated and no gene flow being detected between these individuals (Fig. 2B and Fig. 2C). Compared with the DL and WS-HL populations, 5 YD individuals showed pure YD genetic structures. However, 3 YD individuals had a large proportion of genetic components from the WS-HL population mixed with a small proportion of genetic components from the DL population. These results suggested that some individuals from the WS-HL population inhabit the YD river, and these individuals may have exchanged genes with the DL population. In addition, 4 YD individuals contained equivalent genetic components from the YD and WS-HL populations and showed some admixture with a small proportion of genetic components from the DL population.

**Fig. 2.**
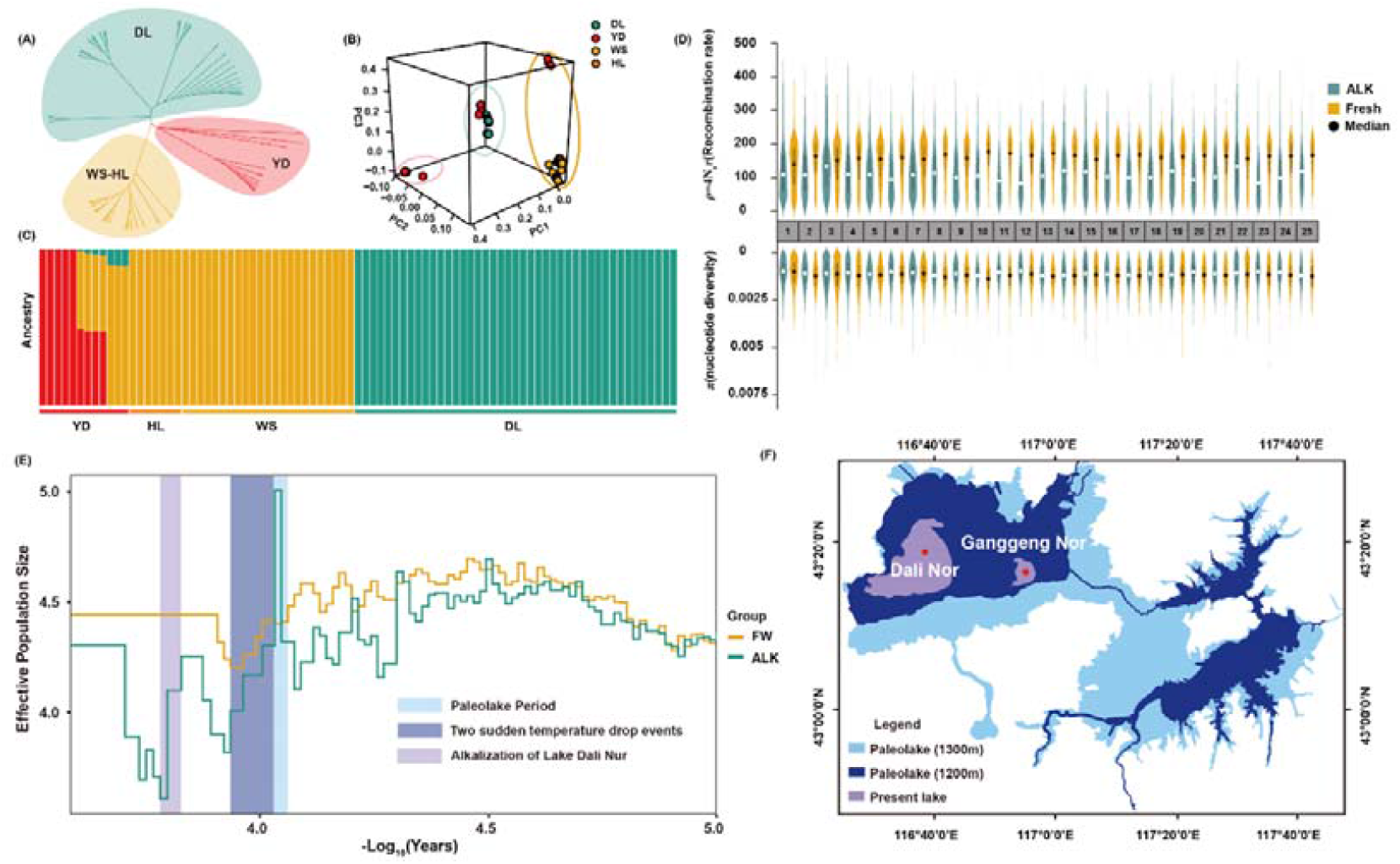
The population genetic and demographic characteristics of the *L. waleckii* ALK and FW populations. (A) Maximum likelihood tree of the relationships between the 85 *L. waleckii* samples based on SNPs. Individuals from different populations are represented by different colours. (B) 3D plot visualizing the principal component analysis (PCA). (C) The structure analysis of 85 *L. waleckii* (K=3) samples. Colours in each column represent the likelihood of an *L. waleckii* individual being assigned to a source population; (D) The recombination and nucleotide diversity statistics for 25 chromosomes between the ALK and FW populations. (E) The effective population size and demographic history of the ALK and FW populations of *L. waleckii*. (F) The schematic diagram of the drainage area of the ancient Paleolake and current Lake Dali Nur.

Natural selection should leave several clear population genetic signatures based on survival in the extremely alkaline environment,. Using the DL population as the alkaline-surviving population (ALK) and WS-HL as the freshwater-inhabiting population (FW), we found that the π values of the ALK population was significantly lower than that of the FW population across the 25 chromosomes (*t test, P*=1.1×10^−4^) (Fig. 2D and Supplymentary File1: Table S29). Statistical analysis also showed that the recombination rate of ALK was significantly lower than that of the FW population (Fig. 2D and Supplymentary File1: Table S29). These results supported the hypothesis that *L. waleckii* may experience an intense selective sweep in the process of adapting to an extreme alkaline environment, which may lead to an independent bottleneck effect of the ALK population compared with the FW population. Hence, we reconstructed the demographic history of the ALK and FW populations. The results showed that approximately 13,000 years ago, the ALK population size dramatically expanded over a thousand years (Fig. 2E). In this period, with the end of the last glacial period, an ancient Lake Dali Nur with a wide drainage area was formed by the convergence of many glacial meltwater layers (Fig. 2F)^42^. With two sudden temperature drop events (Younger Dryas and 8200 BP cold event), the historical populations of both ALK and FW experienced different degrees of decline. In this period, Lake Dali Nur entered a phase of slow contraction^42^. Affected by the monsoon, Lake Dali Nur began to shrink rapidly approximately 6,600 years ago. Because Lake Dali Nur is located inland in Northeast Asia, the warm, humid summer monsoon in East Asia cannot bring precipitation to it. In winter, Lake Dali Nur was controlled by the dryly cold monsoon from the Siberian and Mongolian Plateau (Supplymentary File1: Fig. S10)^23^. Due to the continuous evaporation of water caused by the dry monsoon, the area of Lake Dali Nur has decreased sharply, and water began to alkalize. In this process, the ALK population size dropped and reached its lowest level approximately 6,000 years ago (Fig. 2E). Subsequently, the ALK population gradually adapted to the extreme alkaline environment. Moreover, the ALK population began to recover through opportune occupation of the vacant ecological niche that was caused by the mass extinction of other fishes in Lake Dali Nur in this period. Undoubtedly, the bottleneck effect experienced by the ALK population in the process of adapting to an alkaline environment eventually led to its lower recombination rate and nucleotide diversity compared with the FW population.

### Selective signatures underlying alkaline adaptation in the *L. waleckii* population

Natural selection could leave imprints on specific regions in the genome, such as highly differentiated genetic loci and significant changes in genetic diversity. To identify the candidate genomic regions under selective sweeps in the ALK genome, we scanned the genome-wide variations and allele frequency spectra based on the prospects of approximately 7.0 million SNPs and INDELs. We identified 494 and 488 candidate genes by Fst and π ratios (π_FW/ALK_), respectively; these genes were related to hypoxia tolerance, ion transport, acid-base regulation and nitrogen metabolism (Fig. 3A; Supplymentary File2: Table S30-33). The enrichment categories of 788 candidate genes were associated with oxygen transport (GO:0015671), metal ion binding (GO:0046872), transmembrane transport (GO:0055085), carbonate dehydratase activity (GO:0004089), nitrogen compound transport (GO:0071705), and amino acid transport (GO:0006865), possibly suggesting selection pressure on hypoxia tolerance, ion transport, acid-base regulation and nitrogen metabolism during adaptation to the extreme environment in Lake Dali Nur (Fig. S11, Supplymentary File2: Table S34 and Table S35). In addition, to provide gene expression level evidence linked to the candidate genes, we collected fresh gill, kidney and liver tissue from the ALK and FW populations and analysed RNA-seq data (Supplymentary File1: Table S36). We identified 5,014, 4,848 and 4,468 DEGs in the gill, kidney and liver, respectively (Supplymentary File2: Table S37-Table S45; Supplymentary File1: Fig. S12).

**Fig. 3.**
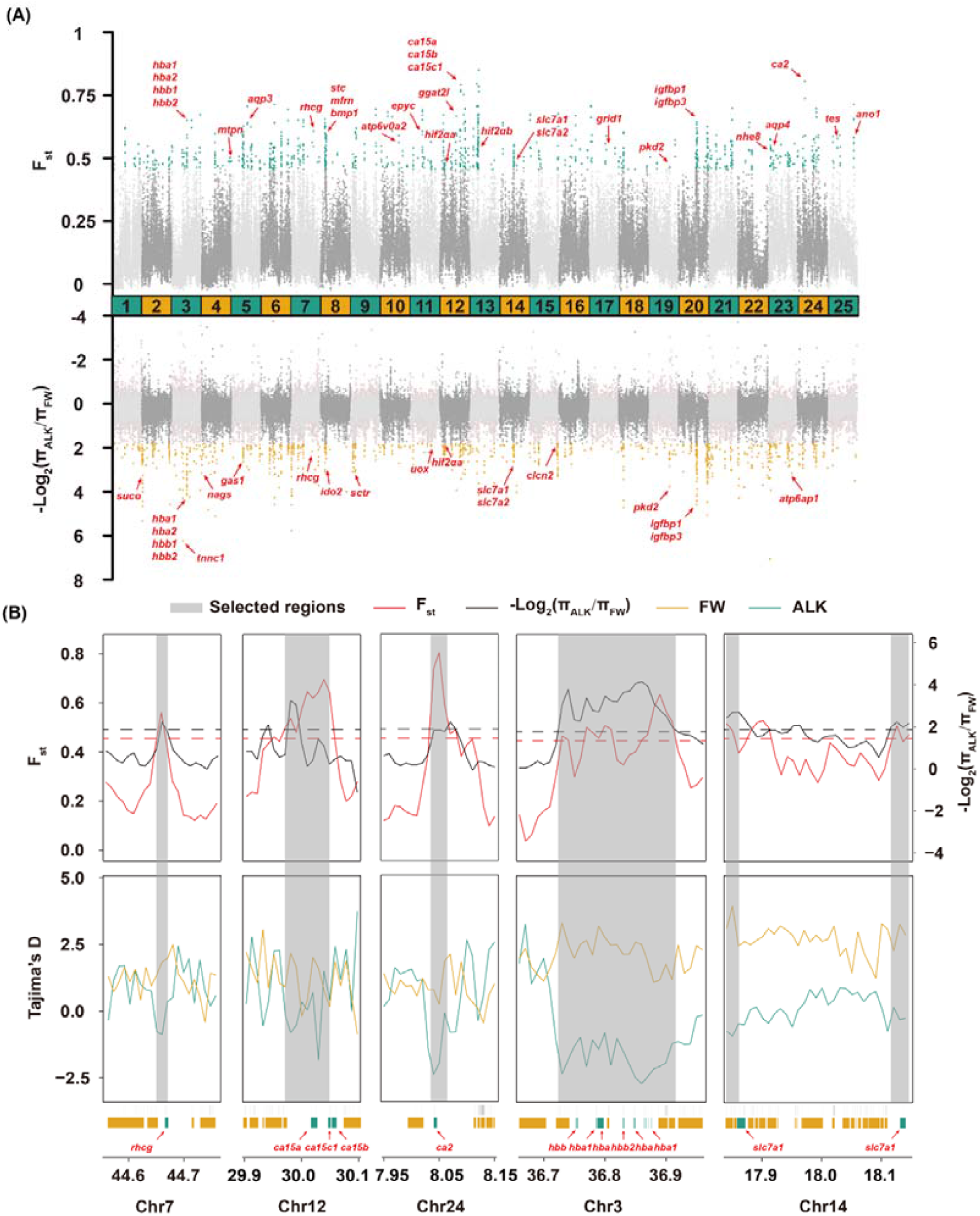
Genomic regions and candidate genes of the selected ALK *L. waleckii* population. (A) Selective scanning of the Fst and π ratio values between the ALK and FW *L. waleckii* populations. The important candidate genes are identified by red arrows. (B) Selective scanning of 13 selected genes. The F_st_, π ratios and Tajima’s D values were plotted using 10,000 bp sliding windows. Genome annotations are shown at the bottom.

Lake Dali Nur is located in the Inner Mongolia Plateau with an average altitude of 1,226 m above sea level and can experience a freezing period of approximately 5 months every year^43^. Therefore, enduring long-term hypoxia is key to the survival of fish in Lake Dali Nur. On chromosome 3 of *L. waleckii*, we identified 200 kb regions, including 12 haemoglobin subunit (*hba*) and 5 haemoglobin subunit beta (*hbb*) genes, that showed a significant selective sweep signal, which was also supported by Tajima’s D analysis (Figure 3B). Among vertebrates, haemoglobin plays a pivotal role in adapting to long-term high-altitude hypoxic environments. For example, several positive selection sites were identified in the Hb genes of Schizothoracinae fishes, and they may accelerate the process of the functional divergence of Hb isoforms^44^. In addition, two copies of the endothelial PAS domain-containing protein 1 (*hif2*_α_) gene, which encodes the transcription factor HIF2α, were detected by Fst and π ratio analysis (Fig. 3B). We also observed elevated expression of *hif1*_α_and *hif2*_α_ in the gills of alkaline-acclimated *L. waleckii* compared with the freshwater population (Supplymentary File2: Table S29). These genes could accelerate erythrocyte synthesis and increase the concentration of haemoglobin in blood^45^. Evidence of natural selection in EPAS1 was found in the Tibetan region, which helped these fish adapt to the extremely low oxygen environment on the Tibetan Plateau^46^. Furthermore, based on RNA sequencing, *hif1*α*B* and *hif2*α*A* might be involved in the high-altitude hypoxia adaptation of *Triplophysa dalaica*^47^. This evidence implied that *hb* and *hif2*α, as the key hypoxia response genes, evolved quickly to adapt to the high-altitude environment.

### Adaptive evolution of carbonic anhydrase in the Lake Dali Nur population

Fish living in Lake Dali Nur need to survive in a continuous carbonate alkaline environment. For most freshwater teleosts, intracellular acid-base regulation occurs by the excretion or uptake of carbon dioxide (CO_2_) and HCO3^-^ through the reversible hydration/dehydration reactions of 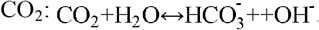. For *L. waleckii*, we identified 3 carbonic anhydrase (CA) genes (*ca15a, ca15c1* and *ca15b*) on chromosome 13in the ALK and FW populations (Fig. 3A and Fig. 3B). In addition, *ca2* was also detected on chromosome 24 and showed a highly differentiated signal (Fst=0.74) (Fig. 3B and Supplymentary File2: Table S31). CAs, as the key zinc metalloenzymes, can catalyse reversible CO_2_ hydration/dehydration reactions^48^. In addition, CAs also provide H^+^ and HCO_3_^-^ for the exchange of Cl^-^/HCO_3_^-^ and Na^+^/H^+^ and Na^+^/HCO_3_^-^ cotransport, which is conducive to maintaining acid-base balance and homeostasis of the internal environment (Fig. 4A)^49^. Among vertebrates, CAs are divided into three groups according to subcellular localization and catalytic activity. (1) Intracellular: CA1, CA2, CA3, CA7, CA13 and CAhz function in the cytoplasm, and CA5 works in the mitochondrion. (2) Extracellular: CA4, CA9, CA12, CA14, and CA15 function in the plasma membrane, and CA6 is secreted into the extracellular matrix. (3) CA-related protein (CA-PR): CA8, CA10 and CA11 lack catalytic activity (Fig. 4B) ^50^. In our genome-wide scan for signatures of selection, four CA genes were identified with significant differentiation signals. Furthermore, we used 18 CAs of zebrafish as queries and identified 19 CAs in *L. waleckii* (Supplymentary File1: Fig. S13, Fig. S14; Supplymentary File2: Table S46, Table S47 and Supplement material 3). In *L. waleckii*, CA15c has two copies, where it only has one copy in zebrafish (Supplymentary File2: Table S48). Compared with the FW population, we observed elevated expression of CA5a, CA9, CA15a, and CA15c1 and decreased expression of CA2 and CAr15 in the gills of the ALK population (Fig. 4C). In the kidney, CAhz was upregulated, and CA2, CA4a, and CA4c were downregulated. In addition, CA2 and Car15 were upregulated, and CA4a and CA8 were downregulated in the liver. To accurately locate the adaptive evolutionary sites of CAs, we calculated the Fst, heterozygosity, and allele frequency of 3,721 SNPs and INDELs in 19 CAs of *L. waleckii* (Fig. 4D and Supplymentary File2: Table S48). Twenty SNPs and 2 INDELs were highly differentiated (Fst > 0.9; MAF_ALK_ < 0.1) between the ALK and FW populations, including 5 nonsynonymous SNPs and 1 synonymous SNP (Supplymentary File2: Table S49). Interestingly, all 22 SNVs were distributed within 3 CA15 copies on chromosome 3. In CA15a, a nonsynonymous SNP was identified in exon 2, which caused an amino acid change from glutamic acid (E) to aspartic acid (D) in the FW *L. waleckii* population (Figure 4E). In CA15c, a nonsynonymous SNP was identified in exon 9. In addition, a SNP mutation was also detected in the 3’ UTR of CA15c. In CA15b, three nonsynonymous SNPs were identified in exons 3, 4 and 5. Comparison with related species showed that these amino acid mutations existed only in the ALK *L. waleckii* population (Fig. 4E). The reconstruction of the 3D model of carbonic anhydrase protein from zebrafish showed that these mutations did not change the spatial structure of the three CA15 copies in alkaline and freshwater *L. waleckii* populations (Fig. 4F). Nevertheless, these alkaline-specific amino acid mutations may change the catalytic activity of their encoded CA15 genes.

**Fig. 4.**
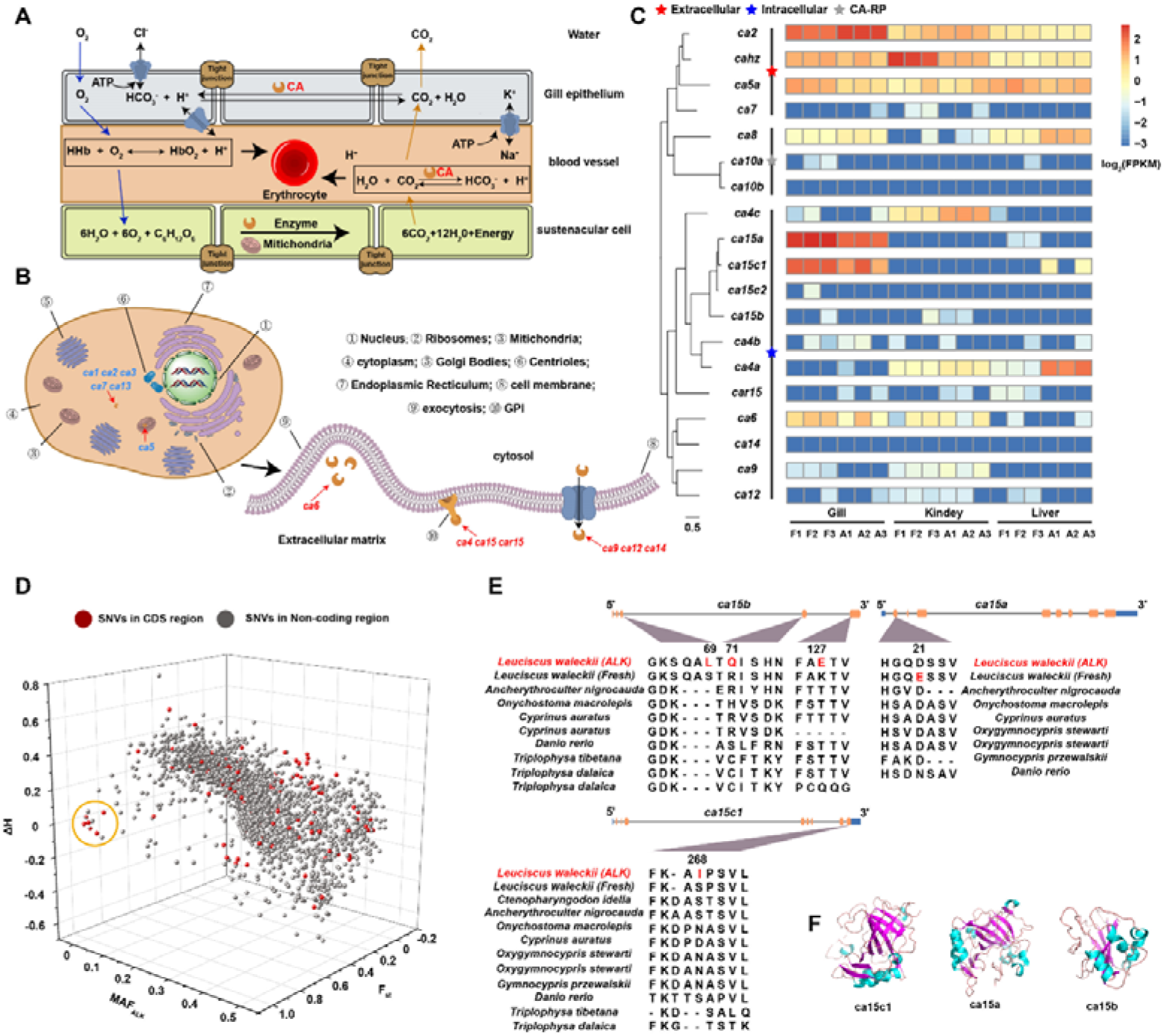
Adaptive evolution of the carbonic anhydrase gene family in *L. waleckii*. (A) The regulatory mechanism of acid-base balance in Cyprinidae fish gills. (B) The distribution of three CA groups according to subcellular localization and catalytic activity. (C) The phylogenetic tree of 19 CA genes in *L. waleckii* and the gene expression heatmap in the gill, liver and kidney between the ALK and FW populations. (D) 3D plot visualizing the highly differentiated SNVs between the ALK and FW populations. The X-axis represents the minimum allele frequency (MAF_ALK_) in the ALK population. The Y-axis represents the Fst between the ALK and FW populations. The Z-axis represents the difference in heterozygosity (ΔH) between the ALK and FW populations. The SNVs located in the CDS region are highlighted by a red dot, and the highly differentiated SNPs are framed with orange circles. (E) The highly differentiated nonsynonymous SNP mutation in 3 copies of CA15 and the protein coding genes of species related to *L. waleckii*. The mutated amino acid is indicated in red. The CDS region is represented by the orange bar, and the UTR is represented by the blue bar. (F) The 3D structure of 3 copies of CA15 in the ALK *L. waleckii* population.

**Fig. 5.**
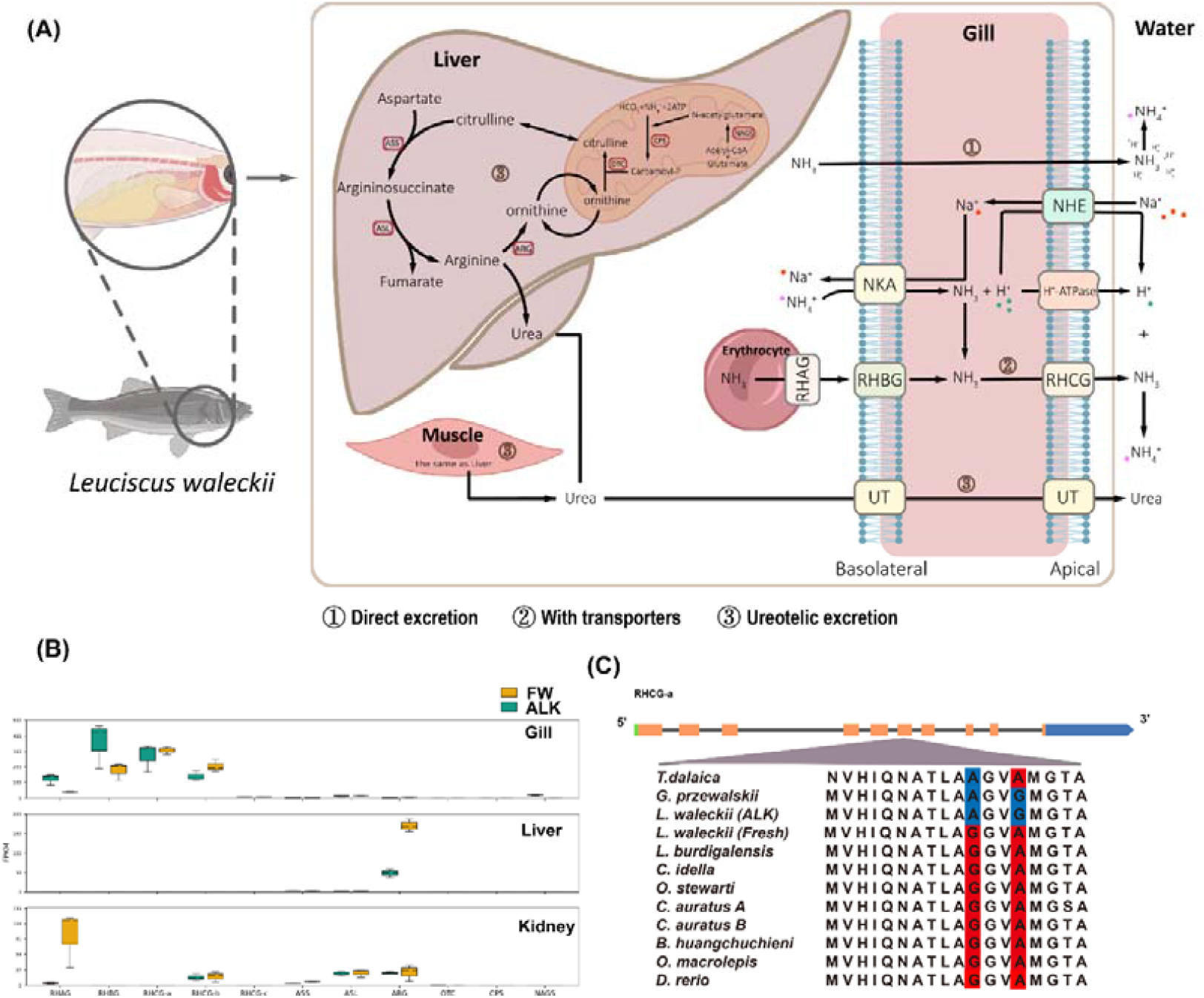
Schematic diagram of the ammonia nitrogen metabolism mechanism in *L. waleckii*. (B) Gene expression heatmap of RH glycoproteins in the gill, liver, and kidney between the ALK and FW populations. (C) The highly differentiated nonsynonymous SNP mutation in RHCG-a and the protein coding genes of species related to *L. waleckii*. The mutated amino acid in the ALK population is indicated in red, and the amino acid in FW is indicated in blue.

In the classical acid-base regulation model in zebrafish, apical H+-ATPase and sodium-hydrogen exchanger 3b (NHE3b) NHE3b provide H^+^ for the CA15-catalysed CO_2_ dehydration reaction, which generates CO_2_ that enters HRCs^12,51^. In contrast, HRCs express cytosolic CA2-like to catalyse CO_2_ hydration and secrete HCO_3_^-^ out of the cell by anion exchange. In Lake Dali Nur, *L. waleckii* upregulated CA15a and CA15c1 to transform excess HCO_3_^-^ to CO_2_. However, to reduce the synthesis rate of HCO_3_^-^ by intracytoplasmic hydration of CO_2_, the expression of CA2 was downregulated to maintain the acid-base balance of the gill, which was in direct contact with the external alkaline environment. Another hypothesis supported that apical membrane-bound CA (CA15a) in the gill could catalyse the CO_2_ hydration and provide the excess protons to NH_3_ to form NH_4_^+52^. Intracellularly, NH_4_^+^ releases H^+^ to form NH_3_, which is transported to the external environment by Rhcga. In these processes, the hydrogen potential difference between the intracellular and extracellular space facilitates Na^+^/H^+^ exchange. However, our expression data for the ALK *L. waleckii* population showed that *slc26a6* was upregulated in the gill, which demonstrates a role of Cl^−^/HCO_3_^−^ exchangers in Cl^-^ uptake (Supplymentary File2: Table S29). Hence, it is more likely that CA15a catalyses CO_2_ dehydration in ALK *L. waleckii* gills, which contributes to the bicarbonate potential difference between the inside and outside of the cell membrane.

### Convergent evolution of *rhcg* in alkaline-adapted Cyprinidae species

Nitrogen metabolism occurs continuously in animals, resulting in the accumulation of toxic ammonia that needs to be excreted or detoxified^17^. For fish living in freshwater, ammonia can easily cross the gills, so it is usually considered to be excreted directly, mainly through Rhesus glycoproteins (Rh)^17^. However, released NH_3_ cannot be trapped by insufficient H^+^ in some aquatic environments, such as alkaline Lake Dali Nur, making it nearly impossible to sustain an ammonia concentration gradient outside the gill^53^. In extremely alkaline conditions where ammonia excretion is inhibited, some fish can slow the build-up of ammonia internally by reducing the ammonia production rate^54^. Moreover, those fish in alkaline water have evolved alternative strategies to eliminate excess ammonia. *Bostrychus sinensis* detoxifies endogenous and exogenous ammonia to neutral glutamine, which participates in various mitochondrial anabolic reactions^55^. *Clarias batrachus*, an air-breathing walking catfish, can increase the content of nonessential free amino acids in the body to respond to the stimulation of hyperammonaemia stress^56^. Moreover, Magadi tilapia produces particularly high rates of urea by the ornithine-urea cycle (OUC) present in muscles as well as in the liver; this urea is subsequently excreted across the gill by a type of urea transporter (*mtUT*)^53^. In the *L. waleckii* ALK population, the ammonium transporter Rh type C (*rhcg-a*) was found within the selected sweep region on chromosome 7 (Figure 3A and Figure 3B). In addition, a series of other genes related to urea synthesis and transport, such as *nags, aqp3* and *aqp4*, showed different degrees of selected signals. Based on RNA-seq, we found that *umod, aqp8* and *aqp10* were upregulated in the kidneys of the ALK population compared with the FW population. The transmembrane transport of arginine (Arg) and ornithine (Orn) guarantees the urea cycle. We identified a selective sweep in two copies of high affinity cationic amino acid transporter 1 (*slc7a1*) on chromosome 14; these genes are involved in the transport of arginine and ornithine (Fig. 3B and Table S42). This evidence once again supports the possibility that *L. waleckii* might excrete urea to avoid the potential accumulation of nitrogenous waste under extreme alkaline environment stress.

To clarify the potential mechanism of ammonia excretion of alkali-adapted *L. waleckii* among such variable pathways, related genes that have already been reported to be associated with excreting ammonia by other researchers were identified and searched across the whole genome^17,53,57-60^. Then, the abundance of the corresponding mRNAs represented the expression of each gene and helped determine which pathway was dominant. Our RNA-seq results implied that for alkali-living *L. waleckii*, the Rh family, especially glycoprotein members, plays the main role in ammonia excretion. Although many related genes have been considered to be rapidly evolving and positively selected genes or their expression is regulated in response to alkaline environments^61^, pervasive molecular adaptation related to ammonia excretion genes has not been previously described. Hence, we identified 7 RH proteins in *L. waleckii* that had 6 shared motifs (Supplymentary File1: Fig. S15, Fig. S16 and Supplymentary File2: Table S51). According to the genome-wide SNP database constructed previously, a total of 764 SNVs from Rh glycoproteins were used to continually explore whether significant differentiation related to these genes exists between freshwater and alkaline water populations (Supplymentary File2: Table S52). Therefore, SNPs with a higher Fst (>0.8) and located within CDS areas were discussed in detail. We still found 5 SNPs under our strict selection criterion (Supplymentary File2: Table S53). In addition, all of these SNPs are nonsynonymous changes that affect the amino acid sequence, and all mutations are in the *rhcga* gene. The products of the *rhcg* genes, of which we found three copies in the genome of *L. waleckii*, were able to move NH_3_ across the apical membrane of the branchial structure^60^. Moreover, all 5 nonsynonymous mutations were counted separately when discussing the conservation of nucleotides at every single population. From our results, the alkaline water population exhibited a more unified sequence for *rhcga*, which suggested that some evolutionary selection pressures had affected this gene^60^. Moreover, all 5 nonsynonymous mutations were counted separately when discussing the conservation of nucleotides at every single population. From our results, the alkaline water population exhibited a more unified sequence for *rhcga*, which suggested that some evolutionary selection pressures had affected this gene.

Based on the smaller population (in particular, a lower recombination rate or lower level of nucleotide diversity in the genome) and adaptation to an extreme alkali environment, we proposed two possible hypotheses to explain the more conserved Rhcga protein sequences in the alkaline papulation: 1. genetic drift and 2. selective sweep. Fortunately, in addition to Amor Ide living in such an alkaline water area, previous studies have determined that other teleost species have evolved some characteristics to survive in the same lake (*Triplophysa dalaica* and *Carassius auratus*) or other similar conditions (*Gymnocypris przewalskii* in Qinghai Lake)^9,62,63^. With sufficient genetic information on these different species living under similar conditions, we checked all the *rhcga* alignments based on sequence similarity and discovered high convergence among four alkaline survivors at two of the seven loci that differentiated freshwater and alkali-water *L. waleckii* populations. At least two alkali-adapted species shared the same amino acid substitution, with some cases sharing identical residues in all four alkaline representatives compared to the completely conserved protein sequence in other teleost fish. The 3D reconstruction of the protein model showed that two mutations did not change the spatial structure of Rhcga in the alkaline and freshwater *L. waleckii* populations (Supplymentary File1: Fig. S17). We then not only confirmed that convergent molecular adaptation to alkaline water arises from selection acting on the same residues in the Rhcga gene but also suggested that common mutations in this gene might play a critical role in the function or structure of ammonia excretion. Furthermore, the convergent substitutions at the same loci in different species that have adapted to similar aquatic conditions implied that the more highly conserved Rhcga sequences in alkaline water populations were likely due to strict sweep selection rather than simple genetic drift.

Although the point at which extremely alkaline survivors possess the capacity for alternative ammonia excretion has long been established, the precise molecular adaptations within genes involved in nitrogen metabolism have not been widely discussed. Our results suggested that the most likely pathway for teleost fish in Lake Dali Nur to excrete fatal ammonia is active ion transport, and we revealed that distant phylogenetic relationships evolved convergent mutations in the same gene to deal with similar environmental pressures. Indeed, the stable genetics of *L. waleckii* in alkaline water were caused by selection under abiotic stress, which enables this species to perform normal physiological activities in such an unusual environment.

## Conclusion

The adaptation of *L. waleckii* to an alkaline lake represents the remarkable adaptability of a species to an alkaline environment. We developed a chromosome-level genome of *L. waleckii* inhabiting an extremely alkaline environment, which provided an important genomic resource for the exploitation of alkaline water fishery resources and adaptive evolution research across teleost fish. Based on comparative genomics, several specific characteristics of adaptive changes in *L. waleckii* regarding gene expansion, transposable elements and selection pressures were detected. Based on the resequencing of 85 *L. waleckii* individuals from divergent populations, genome scans further revealed historical population size fluctuations associated with lacustrine areas and the significant selective sweep regions of Lake Dali Nur *L. waleckii*. These regions harboured a set of candidate genes involved in hypoxia tolerance, ion transport, acid-base regulation and nitrogen metabolism. In particular, several alkali population-specific amino acid mutations were identified in CA15 gene copies. In addition, two convergent evolution amino acid mutation sites were detected in *rhcga* in several alkali environment-adapted Cypriniformes fish.

## Materials and Methods

### Ethics statement

All zoological experiments were carried out according to the recommendations of the Animal Care and Use Committee of Xiamen University (Xiamen, China). *L. waleckii* individuals were euthanized in MS222 (Guangzhou Yibaolai Biochemical Co., Ltd; Guangdong, China) solution before tissue collection.

### Sample collection and genome sequencing

A healthy female *L. waleckii* was collected from Lake Dali Nur, Inner Mongolia (43°22′43′N, 116°39′24′E) (Supplymentary File1: Fig S1b); fresh muscle was immediately frozen in liquid nitrogen for 20 min and then stored at −80 °C for DNA sequencing. The details of DNA extraction and genome sequencing are described in the Appendix Materials and Methods.

### Genome assembly and annotation

To obtain chromosome-level whole genome assembly for *L. waleckii*, we utilized a combined approach of Illumina, PacBio and Hi-C technologies for genome assembly and chromosome-level scaffolding. The strategy of genome assembly was as follows: 1) continuous long-read (CLR) clean reads were assembled using wtdbg2 ^64^ with default parameters. The high-fidelity contig sets were produced by using a combination of circular consensus CLR reads, and Illumina paired-end reads with sufficient overlap to merge into single extended accurate reads; 2) The nonredundant contig sets were reordered and scaffolded using the 3D-DNA pipeline^65^; 3) Scaffolds were fine-tuned and discordant contigs were removed from scaffolds using Juicebox (v. 1.5)^66^. Finally, we obtained a chromosome level reference genome of *L. waleckii* containing linkage group information.

Repetitive sequences of the *L. waleckii* genome were annotated using both homology-based search and de novo methods. Repatscout (v. 1.02)^67^, RepeatModeler (v. 2.0.1)^68^ and LTR_Finder (v. 1.07)^69^ were used to detect repeat sequences in the *L. waleckii* genome. The details of gene structure, function annotation and ncRNA annotation are described in the Appendix Materials and Methods.

Assembly completeness and accuracy were evaluated by multiple methods. First, the Illumina short reads were remapped to the genome using BWA (v. 0.7.17). Then, we used the Benchmarking Universal Single Copy Orthologues (BUSCO) to test the integrality of the final assembly and the lineage dataset was actinopterygii_odb10.

### Evolutionary and comparative genomic analyses

We used the protein coding genes of *Ctenopharyngodon idella, Ancherythroculter nigrocauda* and *L. waleckii* for genomic collinearity analysis by jcvi (v. 1.2.10). Eight species (*A*.*nigrocauda; C*.*idella; L*.*waleckii; Labeo rohita; Onychostoma macrolepis; Danio rerio; Triplophysa tibetana*; *Triplophysa dalaica*) were used in the comparison analysis. RepeatModeler was firstly used to detect repeat sequences in these genomes. TEclass (v. 2.1.3) was used to further annotate unclassified repeats. Then, combined with Repbase (http://www.girinst.org/repbase), a species-specific repeat sequence library was constructed. Finally, RepeatMasker (v. 4.1.0) was utilized to search and classify repeats based on this library.

Single-copy genes in *L. waleckii* and 9 related species (*T. tibetana, T. dalaica, D. rerio, L. rohita, O. macrolepis, C. idella, Anabarilius sgrahami, A. nigrocauda and Oryzias latipes*) were identified based on gene families constructed from protein sequences of all species employing OrthoFinder (v. 2.5.4)^70^ software. Single-copy orthologous proteins were aligned with MUSCLE (v. 3.8.31). A combined continuous ultralong sequence was constructed from all the translated coding DNA alignments for minimum evolution (ME) phylogenetic tree construction using RAxML. The divergence time was estimated using MCMCTREE (PAML package)^71^ based on the molecular clock data of the Timetree database^47^. The expansion and conversion gene families of *L. waleckii* were identified by CAFÉ (v. 4.2).

To perform the dN/dS analysis, we used BLAST to obtain 10,660 reciprocal best hit (RBH) homologues among *A. nigrocauda, C. idella, D. rerio, L. waleckii* and *O. macrolepis* (BLAST E-value cut-off of 1e-5). We employed the software PRANK-MSA (v140110)^72^ with the following parameters, gaprate = 0.025 and gapext = 0.75 to generate coding sequence alignment for each homologous group. To examine the selective constraints on the genes, we estimated the dN/dS ratio (ω) using PAML (v4.4b)^71^. We used three hypotheses: 1) H0, all branches have the same ω; 2) H1, the branch leading to sunfish has a different ω, whereas the other branches have the same ω; and 3) H2, all branches have independent ω. We used likelihood values and degrees of freedom of the three hypotheses to perform a likelihood-ratio test (LRT). We selected genes whose likelihood values of H1 were significantly larger (adjusted LRT p value of < 0.05) than those of H0, and genes whose likelihood values of H2 are not significantly larger than those of H1. Finally, we identified 369 fast-evolving genes with significant false discovery rate (FDR)-corrected p values (<0.05) in *L. waleckii*.

In addition, we also ran branch-site models (model = 2; NSsite = 2) to detect the genes with positively selected sites in *L. waleckii*. For the null hypothesis, we set ‘fix_omega = 1; omega = 1’, whereas for the alternative hypothesis, we set ‘fix_omega = 0; omega = 1.5’ with the tree ‘((*A. nigrocauda*, (*C. idella, L. waleckii #1*)), *O. macrolepis, D. rerio*)’. Using an FDR-corrected LRT p value (adjusted LRT p value) cut-off of 0.05, we identified 131 positively selected genes in *L. waleckii*.

### Differential gene expression analysis

For RNA sequencing, three tissues (gill, kidney and liver) were dissected and collected from three Lake Dali Nur *L. waleckii* individuals and three Wusuli River *L. waleckii* individuals. Total RNA was extracted from the 3 tissues using TRIzol (Invitrogen, Carlsbad, CA, USA). The high-quality RNA samples were sequenced on Illumina HiSeq 2000 platforms according to the manufacturer’s instructions. After sequencing, the adaptor sequences and low-quality reads (quality score ≤ 20) were eliminated to obtain high-quality clean reads using SolexaQA; these reads were then aligned to a reference genome of *L. waleckii* using HISAT2^73^. Stringtie^74^ was used to assemble genes and identify differentially expressed genes (DEGs). We measured the gene expression level based on the fragments per kilobase of exon model per million mapped reads (FPKM). Genes with expression differences fulfilling statistical significance criteria (q-value < P value, P value < 0.05, |log2 (fold change)| > 2) were regarded as differentially expressed genes. To understand the functions of these DEGs, Gene Ontology (GO) functional enrichment and Kyoto Encyclopedia of Genes and Genomes (KEGG) pathway analyses were carried out using OmicShare tools (www.omicshare.com/tools). The threshold for significant enrichment of gene sets was set as p < 0.05.

### Resequencing and population genetic analysis

Twenty-five *L. waleckii* individuals were collected from Lake Dali Nur (DN), Inner Mongolia, 13 individuals were collected from the Wusuli River (WS), 7 individuals were collected from the Hulan River (HL), and 12 individuals were collected from the Yongding River (YD). The sequencing libraries were constructed by TruePrep DNA Library Prep Kit V2 for Illumina (Vazyme Biotec, Nanjing, China). Whole genome resequencing was performed using the Illumina Novo Seq 6000 platform. Paired-end reads from each individual were aligned to the reference genome using the mem algorithm of Burrows–Wheeler Aligner (BWA). GATK v4.0.5.2 was employed to genotype all individuals under standard procedures and preliminarily filter genetic variation sites with the parameters “QD < 5.0 || FS > 35.0 || MQ < 55.0 || SOR > 3.0 || MQRankSum < -12.5 || ReadPosRankSum < -8.0>“^75^. Finally, VCFTOOLS v0.1.06 was used to strictly filter low-quality sites with the parameters “-mac 2 -min-alleles 2 -max-missing-count 2”^76^.

Based on the SNPs and INDELs, a maximum-likelihood tree was constructed by RAxML v8.2.12 with the GTR + I + G model and 1000 bootstrap replicates^77^. The PCA and structure analysis were performed using GCTA v1.26.0 and Admixture v1.3.0 with all SNPs^78,79^. The 3D plots and structure column chart were drawn in R. After excluding YD and HL due to their insufficient sample size, the recent demographic history of the DL and WS populations was inferred by the trend in effective population size (Ne) changes using smc++ with default parameters (mutation rate and recombination of 2e-9, --knots 150, --em-iteration 500)^80^. Each generation was set to 3 years based on the age at sexual maturity of *L. waleckii*.

### Calculation of the recombination rate, π ratio, Fst, and Tajima’s D, and the identification of selective signatures

The recombination rates of ALK and FW were caudated by recombinationCal.pl with a sliding window of 20 kb. To investigate the selection signals for adaptability to extreme alkaline environments in the ALK population, we first scanned the genome using Fst and π ratios with a sliding window size of 20 kb and a step size of 10 kb. The average of the π ratio values for Dali Nur and the FW river 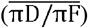 was used to represent the difference in nucleotide diversity. We identified the regions with the 1% highest Fst values (Fst > 0.476) or significant differences in nucleotide diversity 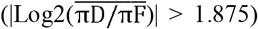. All candidate genes were annotated by blasting candidate regions to the NCBI database. Fine mapping and scanning was performed using Fst, the π ratio and Tajima’s D with a sliding window size of 20 kb and a step size of 10 kb.

### Gene family analysis

All available carbonic anhydrase (CA) and RH protein genes in *H. sapiens, M. musculus, G. gallus, X. tropicalis, D. rerio, G. aculeatus, O. iloticus, T. rubripes, O. latipes, A. nigrocauda, C. auratus, C. idella, O. macrolepis, T. dalaica*, and *T. tibetana* were downloaded from Ensemble (http://asia.ensembl.org/index.html) and the public database GenBank (https://www.ncbi.nlm.nih.gov/genbank/). Furthermore, 20 zebrafish genes were used as queries to search against all available genomics resources by TBLASTN and BLASTP, with an E-value cut-off of 1e-5, to acquire the candidate genes and proteins. The CA and RH proteins were checked using the Pfam (http://pfam.xfam.org/) and Simple Modular Architecture Research Tool (SMART) (http://smart.embl-heidelberg.de/). All CA genes and RH genes were aligned using ClustalW with default parameters. We used the maximum likelihood (ML) method to construct the phylogenetic tree (GTR+G+I model, bootstrap 1000). Then, the nomenclature of all these genes was renamed based on their orthologous genes and their phylogenetic position. The conserved motifs of the CA and RH gene family proteins were analysed using the MEME tool (Multiple Em for Motif Elicitation, (http://meme-suite.org/). The 3D model of selected genes of *L. waleckii* was predicted by SwissModel (https://swissmodel.expasy.org/).

## Author contributions

P. X. and F. P. conceived and designed the project. J. Z., H. B. and Z. Z. collected the samples. Z. Z.,J. Y. and H. L. analyzed the data. Z. Z. and J. Y. wrote the manuscript. T. Z. and P. X. revised the manuscript.

## Competing interests

Authors declare no competing interests.

## Acknowledgments

We thank Heilongjiang Fisheries Research Institute for their help in sample collections. This work was supported by the grants from the National Natural Science Foundation of China (31872561), the National Key R&D Program of China (2019YFE0119000) and the Alliance of International Science Organizations (ANSO-CR-PP-2021-03).

## Data availability

All sequencing data involved in the article have been deposited at the National Genomics Data Center under Bioproject PRJCA009202.

## Notes

### Competing Interest Statement

The authors have declared no competing interest.

## Reference

1. Ma, Z.Y. et al Resequencing a core collection of upland cotton identifies genomic variation and loci influencing fiber quality and yield. Nature Genetics 50, 803-+ (2018).

2. Li, M.Z. et al Genomic analyses identify distinct patterns of selection in domesticated pigs and Tibetan wild boars. Nature Genetics 45, 1431–U180 (2013).

3. Gaither, M.R. et al Genomics of habitat choice and adaptive evolution in a deep-sea fish. Nat Ecol Evol 2, 680–687 (2018).

4. Chen, L. et al The genomic basis for colonizing the freezing Southern Ocean revealed by Antarctic toothfish and Patagonia robalo genomes. GigaScience, 4 (2019).

5. Wolfe, K.H. & Li, W.H. Molecular evolution meets the genomics revolution. Nature Genetics 33, 255–265 (2003).

6. Chen, L., DeVries, A.L. & Cheng, C.H. Convergent evolution of antifreeze glycoproteins in Antarctic notothenioid fish and Arctic cod. Proc Natl Acad Sci U S A 94, 3817–22 (1997).

7. Kaushal, S.S. et al Freshwater salinization syndrome on a continental scale. Proc Natl Acad Sci U S A 115, E574–E583 (2018).

8. Wilkie, M.P., Wood, C.M.J.C.B., Biochemistry, P.P.B. & Biology, M. The adaptations of fish to extremely alkaline environments. 113, 665–673 (1996).

9. Tong, C., Li, M., Tang, Y. & Zhao, K. Genomic Signature of Shifts in Selection and Alkaline Adaptation in Highland Fish. Genome Biol Evol 13(2021).

10. Xu, J. et al Genomic Basis of Adaptive Evolution: The Survival of Amur Ide (Leuciscus waleckii) in an Extremely Alkaline Environment. Mol Biol Evol 34, 145–159 (2017).

11. Xu, J. et al Gene expression changes leading extreme alkaline tolerance in Amur ide (Leuciscus waleckii) inhabiting soda lake. BMC Genomics 14, 682 (2013).

12. Gilmour, K.M. New insights into the many functions of carbonic anhydrase in fish gills. Respiratory physiology neurobiology 184, 223–230 (2012).

13. Galat, D.L., Post, G., Keefe, T.J. & Bouck, G.R. Histological changes in the gill, kidney and liver of Lahontan cutthroat trout, Salmo clarki henshawi, living in lakes of different salinity-alkalinity. Journal of Fish Biology (1985).

14. Goss, G.G., Perry, S.F., Wood, C.M. & Laurent, P. Mechanisms of ion and acid-base regulation at the gills of freshwater fish. Journal of Experimental Zoology 263, 143–159 (2010).

15. Zhang, R.Y. et al Local adaptation of Gymnocypris przewalskii (Cyprinidae) on the Tibetan Plateau. Scientific Reports 5(2015).

16. Wang, Y.S. et al Unusual physiology of scale-less carp, Gymnocypris przewalskii, in Lake Qinghai: a high altitude alkaline saline lake. Comp Biochem Physiol A Mol Integr Physiol 134, 409–21 (2003).

17. Ip, Y.K. & Chew, S.F. Ammonia production, excretion, toxicity, and defense in fish: a review. Front Physiol 1, 134 (2010).

18. Randall, D. et al Urea excretion as a strategy for survival in a fish living in a very alkaline environment. Nature 337, 165–166 (1989).

19. Iwata, K., Kajimura, M. & Sakamoto, T. Functional ureogenesis in the gobiid fish Mugilogobius abei. J Journal of Experimental Biology 203, 3703–3715 (2000).

20. Kavembe, G.D., Franchini, P., Irisarri, I., Machado-Schiaffino, G. & Meyer, A. Genomics of Adaptation to Multiple Concurrent Stresses: Insights from Comparative Transcriptomics of a Cichlid Fish from One of Earth’s Most Extreme Environments, the Hypersaline Soda Lake Magadi in Kenya, East Africa. Journal of Molecular Evolution 81, 90–109 (2015).

21. Wilkie, M.P. & Wood, C.M. The adaptations of fish to extremely alkaline environments. Comparative Biochemistry and Physiology B-Biochemistry & Molecular Biology 113, 665–673 (1996).

22. Zhang, Y. et al Genome evolution trend of common carp (Cyprinus carpio L.) as revealed by the analysis of microsatellite loci in a gynogentic family. Journal of Genetics and Genomics 35, 97–103 (2008).

23. Xiao, J., Si, B., Zhai, D., Itoh, S. & Lomtatidze, Z. Hydrology of Dali Lake in central-eastern Inner Mongolia and Holocene East Asian monsoon variability. Journal of Paleolimnology 40, 519–528 (2008).

24. Wang, S.Y. et al Resequencing and SNP discovery of Amur ide (Leuciscus waleckii) provides insights into local adaptations to extreme environments. Scientific Reports 11(2021).

25. Zhao, X.F. et al Identification and Analysis of Long Non-coding RNAs in Leuciscus waleckii Adapted to Highly Alkaline Conditions. Frontiers in Physiology 12(2021).

26. Wang, B.S. et al Complete mitochondrial genome of Leuciscus waleckii (Cypriniformes: Cyprinidae: Leuciscus). Mitochondrial DNA 24, 126–128 (2013).

27. Xu, J. et al Transcriptome Sequencing and Analysis of Wild Amur Ide (Leuciscus waleckii) Inhabiting an Extreme Alkaline-Saline Lake Reveals Insights into Stress Adaptation. Plos One 8(2013).

28. Wang, X. et al Genomic basis of evolutionary adaptation in a warm-blooded fish. The Innovation 3(2022).

29. Zhou, Z. et al The sequence and de novo assembly of Takifugu bimaculatus genome using PacBio and Hi-C technologies. Sci Data 6, 187 (2019).

30. Xu, P. et al The allotetraploid origin and asymmetrical genome evolution of the common carp Cyprinus carpio. Nat Commun 10, 4625 (2019).

31. Chen, L. et al Chromosome-level genome of Poropuntius huangchuchieni provides a diploid progenitor-like reference genome for the allotetraploid Cyprinus carpio. Mol Ecol Resour 21, 1658–1669 (2021).

32. Wang, Y.P. et al The draft genome of the grass carp (Ctenopharyngodon idellus) provides insights into its evolution and vegetarian adaptation. Nature Genetics 47, 625–631 (2015).

33. Jian, J.B. et al Whole genome sequencing of silver carp (Hypophthalmichthys molitrix) and bighead carp (Hypophthalmichthys nobilis) provide novel insights into their evolution and speciation. Molecular Ecology Resources 21, 912–923 (2021).

34. Mulch, A. & Chamberlain, C.P. Earth science - The rise and growth of Tibet. Nature 439, 670–671 (2006).

35. Schrader, L. & Schmitz, J. The impact of transposable elements in adaptive evolution. Mol Ecol 28, 1537–1549 (2019).

36. Balen, J., Demeulemeester, A.A., Frölich, M., Mohrmann, K. & Souverijn, J. Gamma-glutamyltransferase, (San Memoboek, 2012).

37. Dafonsecawollheim, F. Deamidation of Glutamine by Increased Plasma Gamma-Glutamyl-Transferase Is a Source of Rapid Ammonia Formation in Blood and Plasma Specimens. Clinical Chemistry 36, 1479–1482 (1990).

38. Heisterkamp, N., Groffen, J., Warburton, D. & Sneddon, T.P. The human gamma-glutamyltransferase gene family. Human Genetics 123, 321–332 (2008).

39. Zhang, N.S., Li, H.Y., Liu, J.S. & Yang, W.D. Gene expression profiles in zebrafish (Danio rerio) liver after acute exposure to okadaic acid. Environmental Toxicology and Pharmacology 37, 791–802 (2014).

40. Murray, B.W., Busby, E.R., Mommsen, T.P. & Wright, P.A. Evolution of glutamine synthetase in vertebrates: multiple glutamine synthetase genes expressed in rainbow trout (Oncorhynchus mykiss). Journal of Experimental Biology 206, 1511–1521 (2003).

41. Supek, F., Bosnjak, M., Skunca, N. & Smuc, T. REVIGO Summarizes and Visualizes Long Lists of Gene Ontology Terms. Plos One 6(2011).

42. Lan, Y.H., Tian, M.Z., Zhang, X.J., Wen, X.F. & Kang, C.J. Evolution of a Late Pleistocene palaeolake in Dali Nor area of southeastern Inner Mongolia Plateau, China. Geoscience Frontiers 9, 223–237 (2018).

43. Ma, C.X. et al Spatial and temporal variation of phytoplankton functional groups in extremely alkaline Dali Nur Lake, North China. Journal of Freshwater Ecology 34, 91–105 (2019).

44. Lei, Y. et al Hb adaptation to hypoxia in high-altitude fishes: Fresh evidence from schizothoracinae fishes in the Qinghai-Tibetan Plateau. International Journal of Biological Macromolecules 185, 471–484 (2021).

45. Tian, H., McKnight, S.L. & Russell, D.W. Endothelial PAS domain protein 1 (EPAS1), a transcription factor selectively expressed in endothelial cells. Genes & Development 11, 72–82 (1997).

46. Beall, C.M. et al Natural selection on EPAS1 (HIF2alpha) associated with low hemoglobin concentration in Tibetan highlanders. Proc Natl Acad Sci U S A 107, 11459–64 (2010).

47. Wang, Y., Yang, L., Wu, B., Song, Z. & He, S. Transcriptome analysis of the plateau fish (Triplophysa dalaica): Implications for adaptation to hypoxia in fishes. Gene 565, 211–20 (2015).

48. Gilmour, K.M. & Perry, S.F. Carbonic anhydrase and acid-base regulation in fish. Journal of Experimental Biology 212, 1647–1661 (2009).

49. Henry, R.P. Multiple roles of carbonic anhydrase in cellular transport and metabolism. Annual Review of Physiology 58, 523–538 (1996).

50. Ferreira-Martins, D. et al A cytosolic carbonic anhydrase molecular switch occurs in the gills of metamorphic sea lamprey. Sci Rep 6, 33954 (2016).

51. Lin, C.H., Shih, T.H., Liu, S.T., Hsu, H.H. & Hwang, P.P. Cortisol Regulates Acid Secretion of H(+)-ATPase-rich Ionocytes in Zebrafish (Danio rerio) Embryos. Front Physiol 6, 328 (2015).

52. Wright, P.A. & Wood, C.M. A new paradigm for ammonia excretion in aquatic animals: role of Rhesus (Rh) glycoproteins. J Exp Biol 212, 2303–12 (2009).

53. Wood, C.M. et al Rh proteins and NH4(+)-activated Na+-ATPase in the Magadi tilapia (Alcolapia grahami), a 100% ureotelic teleost fish. J Exp Biol 216, 2998–3007 (2013).

54. Lim, C.B., Chew, S.F., Anderson, P.M. & Ip, Y.K. Reduction in the rates of protein and amino acid catabolism to slow down the accumulation of endogenous ammonia: a strategy potentially adopted by mudskippers (Periophthalmodon schlosseri snd Boleophthalmus boddaerti) during aerial exposure in constant darkness. J Exp Biol 204, 1605–14 (2001).

55. Peh, W.Y.X., Chew, S.F., Wilson, J.M. & Ip, Y.K. Branchial and intestinal osmoregulatory acclimation in the four-eyed sleeper, Bostrychus sinensis (LacepSde), exposed to seawater. Marine Biology 156, 1751–1764 (2009).

56. Saha, N., Dutta, S. & Haussinger, D. Changes in free amino acid synthesis in the perfused liver of an air-breathing walking catfish, Clarias batrachus infused with ammonium chloride: a strategy to adapt under hyperammonia stress. J Exp Zool 286, 13–23 (2000).

57. Biver, S. et al A role for Rhesus factor Rhcg in renal ammonium excretion and male fertility. Nature 456, 339–43 (2008).

58. Braun, M.H., Steele, S.L., Ekker, M. & Perry, S.F. Nitrogen excretion in developing zebrafish (Danio rerio): a role for Rh proteins and urea transporters. Am J Physiol Renal Physiol 296, F994–F1005 (2009).

59. Ip, A.Y. & Chew, S.F. Ammonia production, excretion, toxicity, and defense in fish: a review. Frontiers in physiology 1, 134 (2010).

60. Wright, P.A. & Wood, C.M. A new paradigm for ammonia excretion in aquatic animals: role of Rhesus (Rh) glycoproteins. Journal of Experimental Biology 212, 2303–2312 (2009).

61. Tong, C., Li, M., Tang, Y. & Zhao, K. Genomic signature of ongoing alkaline adaptation in a Schizothoracine fish (Cyprinidae) inhabiting soda lake on the Tibetan Plateau. bioRxiv, 813501 (2019).

62. Zhou, C. et al The chromosome-level genome of Triplophysa dalaica (Cypriniformes: Cobitidae) provides insights into its survival in extremely alkaline environment. Genome biology and evolution 13, evab153 (2021).

63. Luo, H. et al Full-length transcript sequencing accelerates the transcriptome research of Gymnocypris namensis, an iconic fish of the Tibetan Plateau. Scientific reports 10, 1–11 (2020).

64. Ruan, J. & Li, H. Fast and accurate long-read assembly with wtdbg2. Nat Methods 17, 155–158 (2020).

65. Dudchenko, O. et al De novo assembly of the Aedes aegypti genome using Hi-C yields chromosome-length scaffolds. Science 356, 92–95 (2017).

66. Robinson, J.T. et al Juicebox.js Provides a Cloud-Based Visualization System for Hi-C Data. Cell Syst 6, 256–258 e1 (2018).

67. Price, A.L., Jones, N.C. & Pevzner, P.A. De novo identification of repeat families in large genomes. Bioinformatics 21 Suppl 1, i351–8 (2005).

68. Tarailo-Graovac, M. & Chen, N. Using RepeatMasker to identify repetitive elements in genomic sequences. Curr Protoc Bioinformatics Chapter 4, Unit 4 10 (2009).

69. Xu, Z. & Wang, H. LTR_FINDER: an efficient tool for the prediction of full-length LTR retrotransposons. Nucleic Acids Res 35, W265–8 (2007).

70. Emms, D.M. & Kelly, S. OrthoFinder: phylogenetic orthology inference for comparative genomics. Genome Biol 20, 238 (2019).

71. Yang, Z. PAML: a program package for phylogenetic analysis by maximum likelihood. Comput Appl Biosci 13, 555–6 (1997).

72. Loytynoja, A. Phylogeny-aware alignment with PRANK. Methods Mol Biol 1079, 155–70 (2014).

73. Kim, D., Langmead, B. & Salzberg, S.L. HISAT: a fast spliced aligner with low memory requirements. Nat Methods 12, 357–60 (2015).

74. Pertea, M., Kim, D., Pertea, G.M., Leek, J.T. & Salzberg, S.L. Transcript-level expression analysis of RNA-seq experiments with HISAT, StringTie and Ballgown. Nat Protoc 11, 1650–67 (2016).

75. McKenna, A. et al The Genome Analysis Toolkit: a MapReduce framework for analyzing next-generation DNA sequencing data. Genome Res 20, 1297–303 (2010).

76. Danecek, P. et al The variant call format and VCFtools. Bioinformatics 27, 2156–8 (2011).

77. Stamatakis, A. RAxML version 8: a tool for phylogenetic analysis and post-analysis of large phylogenies. Bioinformatics 30, 1312–3 (2014).

78. Yang, J., Lee, S.H., Goddard, M.E. & Visscher, P.M. GCTA: a tool for genome-wide complex trait analysis. Am J Hum Genet 88, 76–82 (2011).

79. Alexander, D.H., Novembre, J. & Lange, K. Fast model-based estimation of ancestry in unrelated individuals. Genome Res 19, 1655–64 (2009).

80. Patton, A.H. et al Contemporary Demographic Reconstruction Methods Are Robust to Genome Assembly Quality: A Case Study in Tasmanian Devils. Mol Biol Evol 36, 2906–2921 (2019).

