## Supplementary material for "Genomic insights into the adaptive and convergent evolution of *Leuciscus waleckii* inhabiting extremely alkaline environments": Supplymentary File1

**The chromosome-level genome of *Leuciscus waleckii* provide insight into the adaptive and convergent evolution under the extreme alkaline environment in Lake Dali Nur**

Zhixiong Zhou^1,2,3^, Junyi Yang^1,2,3^, Hongzao Lv^1,2,3^, Tao Zhou^1,2,3^, Ji Zhao^1,2,3^, Huaqiang Bai^1,2,3^, Fei Pu^1,2,3^, Peng Xu^1.2.3^

1 State Key Laboratory of Marine Environmental Science, College of Ocean and Earth Sciences, Xiamen University, Xiamen, 361102, China

2 Fujian Key Laboratory of Genetics and Breeding of Marine Organisms, College of Ocean and Earth Sciences, Xiamen University, Xiamen 361102, China

3 Laboratory for Marine Biology and Biotechnology, Pilot National Laboratory for Marine Science and Technology, Qingdao, 266071, China

**SUPPLEMENTARY INFORMATION**

### Materials and Methods

#### Sample collection and sequencing

A healthy female *L. waleckii* was collected from Lake Dali Nur, Inner Mongolia (43°22′43′N, 116°39′24′E) (Fig. S1b); fresh muscle and was immediately frozen in liquid nitrogen for 20 min and then stored at −80 °C for DNA sequencing. The high-molecular-weight (HMW) genomic DNA (gDNA) for PacBio and Hi-C was extracted by AMPure XP beads (Beckman Coulter, High Wycombe, UK) under previous method ^1^. Meanwhile, normal-molecular-weight (NMW) gDNA for Illumina was extracted using the PureLink Genomic DNA Mini Kit (hermo Fisher Scientific, Shanghai, China). Nucleic acid concentrations were quantified by the Qubit fluorometer 4.0 (Thermo Fisher Scientific, Waltham, MA), and integrity was checked by Agilent2100 Bioanalyzer (Agilent Technologies, Palo Alto, Calif). According to Illumina standard operating procedures, a shotgun library with 350 bp insert size was constructed. Illumina raw reads were generated from Hiseq-X Ten platform, after which uncertain bases (represented by “N”) and low-quality reads (Q < 5) were trimmed by SolexaQA++ ^2^. Finally, all clean reads were retained for the evaluation of genome size and polishing of preliminary contigs. The SMRTbell large fragment sequencing library was constructed according to standard procedures provided by Pacific Biosciences and sequenced on PacBio Sequel Ⅱ platform. *Mbol* restriction enzyme was used to digest the HMW gDNA after fixing conformation of HMW gDNA by formaldehyde, followed by repair the 5′ overhangs with biotinylated residues. The isolated DNA was reverse-crosslinked, purified and filtered for biotin-containing fragments after the ligation of blunt-end in situ. Thereafter, the DNA was sheared into fragments by ultrasonic, the sheared DNA ends was subsequently repaired by T4 DNA polymerase, T4 polynucleotide kinase and Klenow DNA polymerase. Then, a dATP was attached to the 3' ends of the end-repaired DNA and 300-500 bp fragments was retrieved by Caliper LabChip Xte (PerkinElmer, USA). The DNA concentration was determined by Qubit and the Illumina Paired End adapters was ligated to the DNA by T4 DNA Ligase. Thereafter, 12 cycles PCR reactions was conducted and products were purified by AMpureXP beads. Finally, sequencing of Hi-C library was performed on Illumina Hiseq X-Ten platform and yielded a total of 102.93 Gb pair-end raw reads and 100.36 Gb was retained after quality control (Table S1).

#### Genome assembly and annotation

Reads from the three types of libraries were used in different assembly stages separately. Illumina sequencing data were used for genome survey, PacBio sequencing data were served in contig assembly, and Hi-C reads were used in chromosome-level scaffolding. In genome survey, the 17 mers frequency of 73 Gb Illumina clean data was counted with 1 bp sliding window by Jellyfish (Table S1 and Figure S2) ^3^. Finally, the proportion of heterozygosity in *L. waleckii* genome was evaluated as 0.56%, and the genome size was estimated as 1125.03 Mb, with a repeat content of 57.61% (Table S2). A total of 62 Gb raw reads were generated by PacBio with a mean insert size of 35,943, resulting in ~54.87X coverage of the k-mer estimated genome size of *L.waleckii* (Tables S3). Long reads generated from the PacBio SEQUEL platform were subsequently processed by a self-correction of errors using Canu^4^. Based on Overlap-Layout-Consensus algorithm, we detected overlaps from input reads and assembled the final String Graph by wtdbg2 ^5^. Subsequently we used FALCON-unzip pipeline to generate phased contig sequences for further calling highly accurate consensus sequences using variantCaller in GenomicConsensus package, which was employed arrow algorithm, and polishing the contigs using Illumina reads by pilon^6^. Finally, we obtained the assembled genome of *L. waleckii* including 6,407 contigs, with the total length and contig N50 length were 1,103.97 Mb and 1.52 Mb respectively (Table S4).

For chromosome-level scaffolding, we first filtered Hi-C reads with the same protocol as Illumina reads. Subsequently, we mapped the Hi-C clean reads to the *de novo* assembled contigs by BWA^7^ with default parameter. We removed the reads that unmapped within 500bp of a restriction enzyme site. Using 3D-DNA, we assembled the chromosome-level scaffolding based on the signal of genomic proximity in Hi-C data sets. In this stage, all parameters were default. As a result, we generated 25 chromosome-level scaffolds with a total length of 1020.34 Mb (92.32% of total length of all contigs), and the chromosomes lengths ranging from 71.37 Mb to 28.42 Mb (Table S9).

Repetitive sequences of the *L. waleckii* genome were annotated using both homology-based search and de novo methods. Combined with Repbase (v. 20,181,026; http://www.girinst.org/repbase), a repeat sequence library was constructed. RepeatMasker (v. 4.1.0) were utilized to search and classify repeats based on this library. TEclass (v. 2.1.3)^8^ was used to further annotate unclassified repeats. The built-in script buildSummary.pl from RepeatMasker (v. 4.1.0) was used to summarize Transposable Elements (TEs) annotation results. Then two scripts, calcDivergenceFromalign.pl and createRepeatLandscape.pl, were used to calculate the Kimura divergence value and draw repeated landscapes. The nucleotide distances between all copies of each TE measured using the Kimura two-parameter method were compared to estimate insertion age ^9^. Tandem Repeats Finder (v. 4. 09)^10^ was used to identify tandem repeats. All repetitive regions except tandem repeats were soft-masked for protein-coding gene annotation. For gene structure prediction, we used both homology-based and de novo strategies to predict genes in the *L. waleckii* genome. For homology-based prediction, we mapped the protein sequences of *Cyprinus carpio^11^*, *Danio rerio*^12^, *Ctenopharyngodon idellus*^13^, *Sinocyclocheilus grahami*^14^ and *L. waleckii*^15^ onto the generated assembly using BLAT^16^ (version 35) with an e-value ≤ 1e-5. Then, we used GeneWise^17^ (version 2.2.0) to align the homologous in the *L. waleckii* genome against the other five teleosts for gene structure prediction. In the de novo approach, we used several software packages, including Augustus (version 2.5.5)^18^, GlimmerHMM (version 3.0.1)^19^, SNAP (version 1.0)^20^, Geneid (version 1.4.4)^21^ and GenScan (version 1.0)^22^. In addition, we also used RNA-seq data to predict the structure of transcribed genes using TopHat (version 1.2)^23^ and Cufflinks (version 2.2.1)^24^. Using EvidenceModeler (version 1.1.0) ^25^, we combined the setof predicted genes generated from the three approaches into a non-redundant gene set and then used PASA (version 2.0.2)^26^ to annotate the gene structures. For gene function annotation, we used BLASTP to align the candidate sequences to the NCBI and Swissport protein databases with E values < 1 × 10−5. Then, we performed the functional classification of GO categories with the InterProScan program (version 5.26)^27^ and used KEGG Automatic Annotation Server (KAAS)^28^ to conduct the KEGG pathway annotation analysis. The programs tRNAScan-SE (v. 1.3.1) and RNAmmer (v. 1.2) were used to predict tRNA and rRNA, respectively. The other ncRNAs were identified by searching against the Rfam database (<http://eggnogdb.embl.de/>).

### Reference

1.	Zhou, Z.X. *et al.* The sequence and de novo assembly of Takifugu bimaculatus genome using PacBio and Hi-C technologies. *Scientific Data* **6**(2019).

2.	Cox, M.P., Peterson, D.A. & Biggs, P.J. SolexaQA: At-a-glance quality assessment of Illumina second-generation sequencing data. *BMC Bioinformatics* **11**, 485 (2010).

3.	Xu, P. *et al.* Genome sequence and genetic diversity of the common carp, Cyprinus carpio. *Nat Genet* **46**, 1212-9 (2014).

4.	Koren, S. *et al.* Canu: scalable and accurate long-read assembly via adaptive k-mer weighting and repeat separation. *Genome Research* **27**, 722-736 (2017).

5.	Myers, E.W. The fragment assembly string graph. *Bioinformatics* **21 Suppl 2**, ii79-85 (2005).

6.	Walker, B.J. *et al.* Pilon: an integrated tool for comprehensive microbial variant detection and genome assembly improvement. *PLoS One* **9**, e112963 (2014).

7.	Li, H. & Durbin, R. Fast and accurate short read alignment with Burrows-Wheeler transform. *Bioinformatics* **25**, 1754-60 (2009).

8.	Abrusan, G., Grundmann, N., DeMester, L. & Makalowski, W. TEclass--a tool for automated classification of unknown eukaryotic transposable elements. *Bioinformatics* **25**, 1329-30 (2009).

9.	Schemberger, M.O. *et al.* DNA transposon invasion and microsatellite accumulation guide W chromosome differentiation in a Neotropical fish genome. *Chromosoma* **128**, 547-560 (2019).

10.	Benson, G. Tandem repeats finder: a program to analyze DNA sequences. *Nucleic Acids Research* **27**, 573-580 (1999).

11.	Xu, P. *et al.* The allotetraploid origin and asymmetrical genome evolution of the common carp Cyprinus carpio. *Nat Commun* **10**, 4625 (2019).

12.	Howe, K. *et al.* The zebrafish reference genome sequence and its relationship to the human genome (vol 496, pg 498, 2013). *Nature* **505**, 248-248 (2014).

13.	Wang, Y.P. *et al.* The draft genome of the grass carp (Ctenopharyngodon idellus) provides insights into its evolution and vegetarian adaptation. *Nature Genetics* **47**, 625-631 (2015).

14.	Yang, J.X. *et al.* The Sinocyclocheilus cavefish genome provides insights into cave adaptation. *Bmc Biology* **14**(2016).

15.	Xu, J. *et al.* Genomic Basis of Adaptive Evolution: The Survival of Amur Ide (Leuciscus waleckii) in an Extremely Alkaline Environment. *Molecular Biology and Evolution* **34**, 145-159 (2017).

16.	Kent, W.J. BLAT - The BLAST-like alignment tool. *Genome Research* **12**, 656-664 (2002).

17.	Birney, E., Clamp, M. & Durbin, R. GeneWise and Genomewise. *Genome Res* **14**, 988-95 (2004).

18.	Stanke, M. & Morgenstern, B. AUGUSTUS: a web server for gene prediction in eukaryotes that allows user-defined constraints. *Nucleic Acids Research* **33**, W465-W467 (2005).

19.	Majoros, W.H., Pertea, M. & Salzberg, S.L. TigrScan and GlimmerHMM: two open source ab initio eukaryotic gene-finders. *Bioinformatics* **20**, 2878-9 (2004).

20.	Korf, I. Gene finding in novel genomes. *BMC Bioinformatics* **5**, 59 (2004).

21.	Parra, G., Blanco, E. & Guigo, R. GeneID in Drosophila. *Genome Res* **10**, 511-5 (2000).

22.	Burge, C. & Karlin, S. Prediction of complete gene structures in human genomic DNA. *J Mol Biol* **268**, 78-94 (1997).

23.	Trapnell, C., Pachter, L. & Salzberg, S.L. TopHat: discovering splice junctions with RNA-Seq. *Bioinformatics* **25**, 1105-11 (2009).

24.	Trapnell, C. *et al.* Transcript assembly and quantification by RNA-Seq reveals unannotated transcripts and isoform switching during cell differentiation. *Nat Biotechnol* **28**, 511-5 (2010).

25.	Haas, B.J. *et al.* Automated eukaryotic gene structure annotation using EVidenceModeler and the program to assemble spliced alignments. *Genome Biology* **9**(2008).

26.	Haas, B.J. *et al.* Improving the Arabidopsis genome annotation using maximal transcript alignment assemblies. *Nucleic Acids Research* **31**, 5654-5666 (2003).

27.	Jones, P. *et al.* InterProScan 5: genome-scale protein function classification. *Bioinformatics* **30**, 1236-40 (2014).

28.	Moriya, Y., Itoh, M., Okuda, S., Yoshizawa, A.C. & Kanehisa, M. KAAS: an automatic genome annotation and pathway reconstruction server. *Nucleic Acids Res* **35**, W182-5 (2007).

29.	Kim, D., Langmead, B. & Salzberg, S.L. HISAT: a fast spliced aligner with low memory requirements. *Nat Methods* **12**, 357-60 (2015).

### Supplementary Figure

#### Fig S1. Sampling location of *L. waleckii*

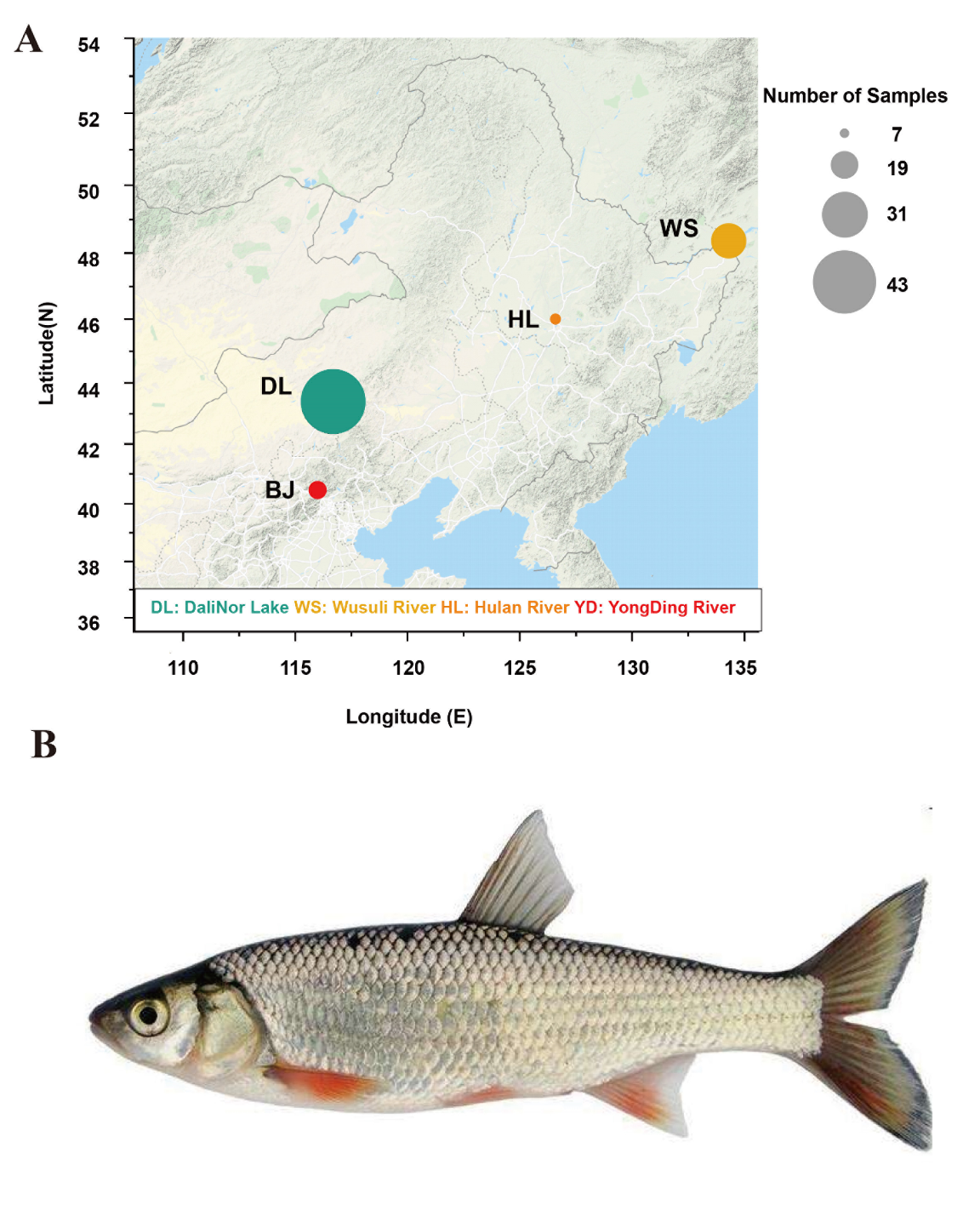

(A) geographic locations of sample collection. (B) The photo of *L. waleckii*.

#### Fig S2. Genome size estimation.

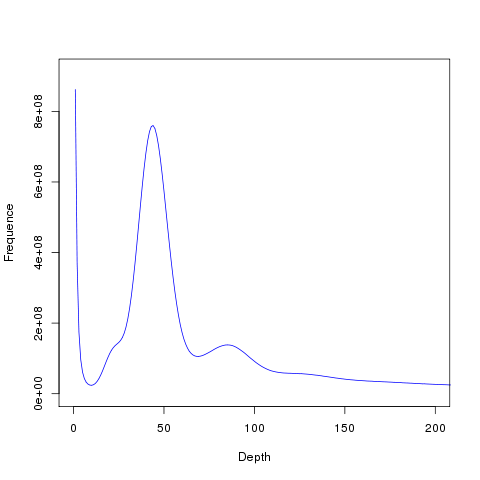

We used 3.7 ×10^9^ sequence reads and obtained 49 × 10^9^ 17-mers. The peak depth is 43. The genome size (G) is correlated with the 17-mer number (N) and the peak of 17-mer frequency (D). Their relationship can be expressed in an empiric formula: G = N / D. The estimated genome size is 1125.03 Mb.

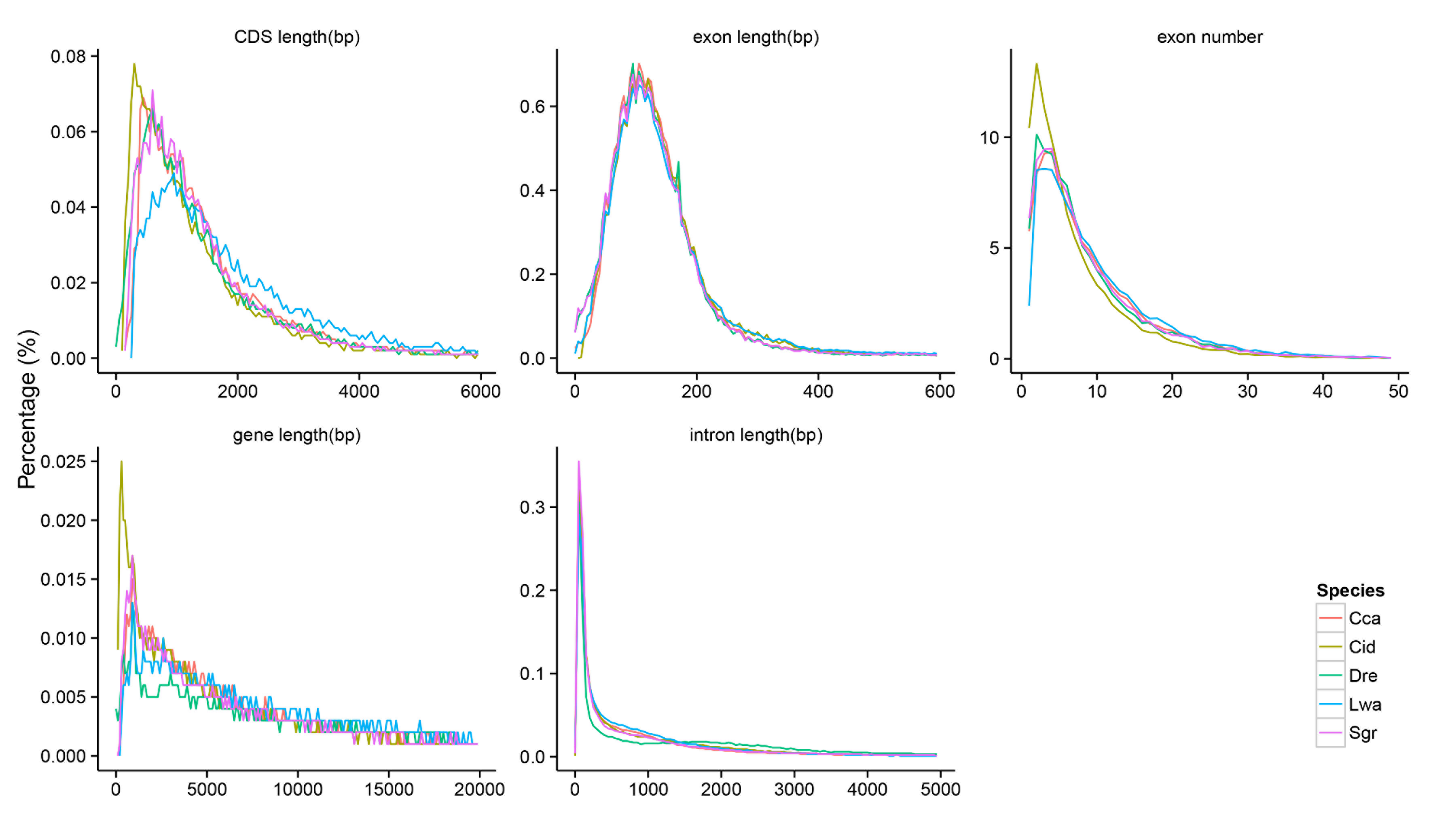

#### Fig S3. The comparison of gene structure elements between Amur Ide and related species.

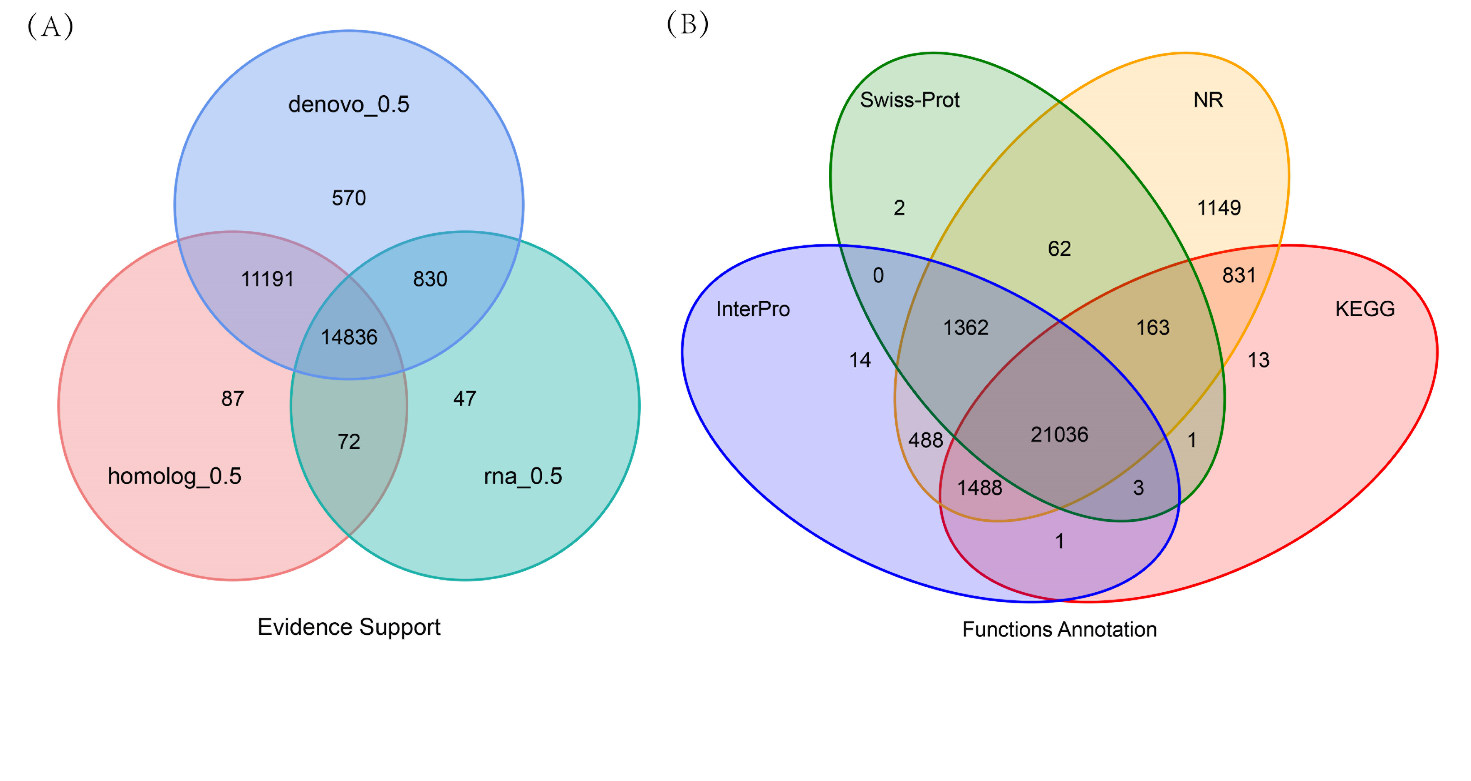

#### Fig S4. Gene structure and function annotation.

(A) Venn diagram of the number of genes with structure prediction based on different strategies. (B) Venn diagram of the number of functionally annotated genes based on different public databases.

#### Fig S5. The Hi-C heatmap of Amur Ide represented the assembly of the chromosome.

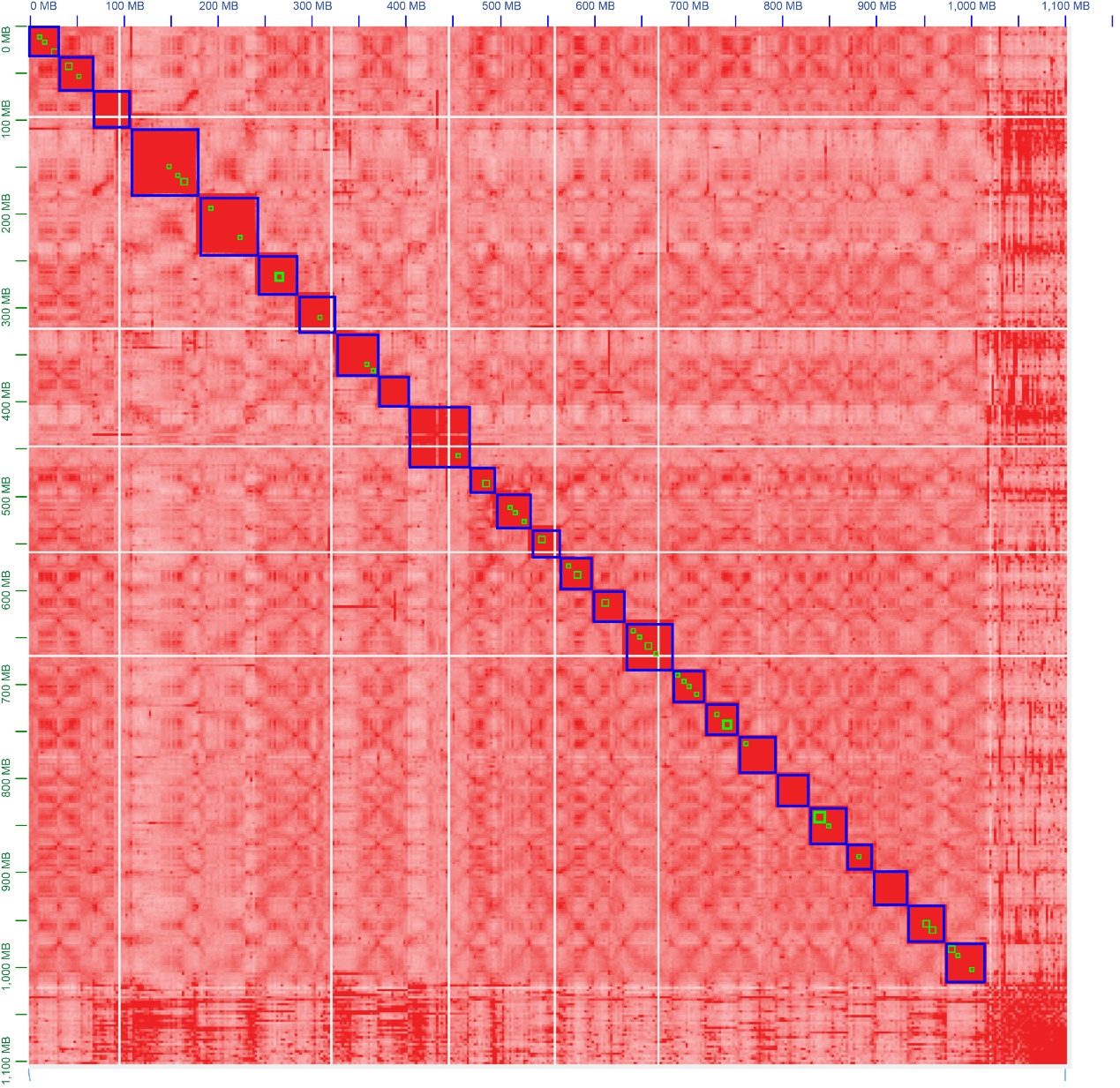

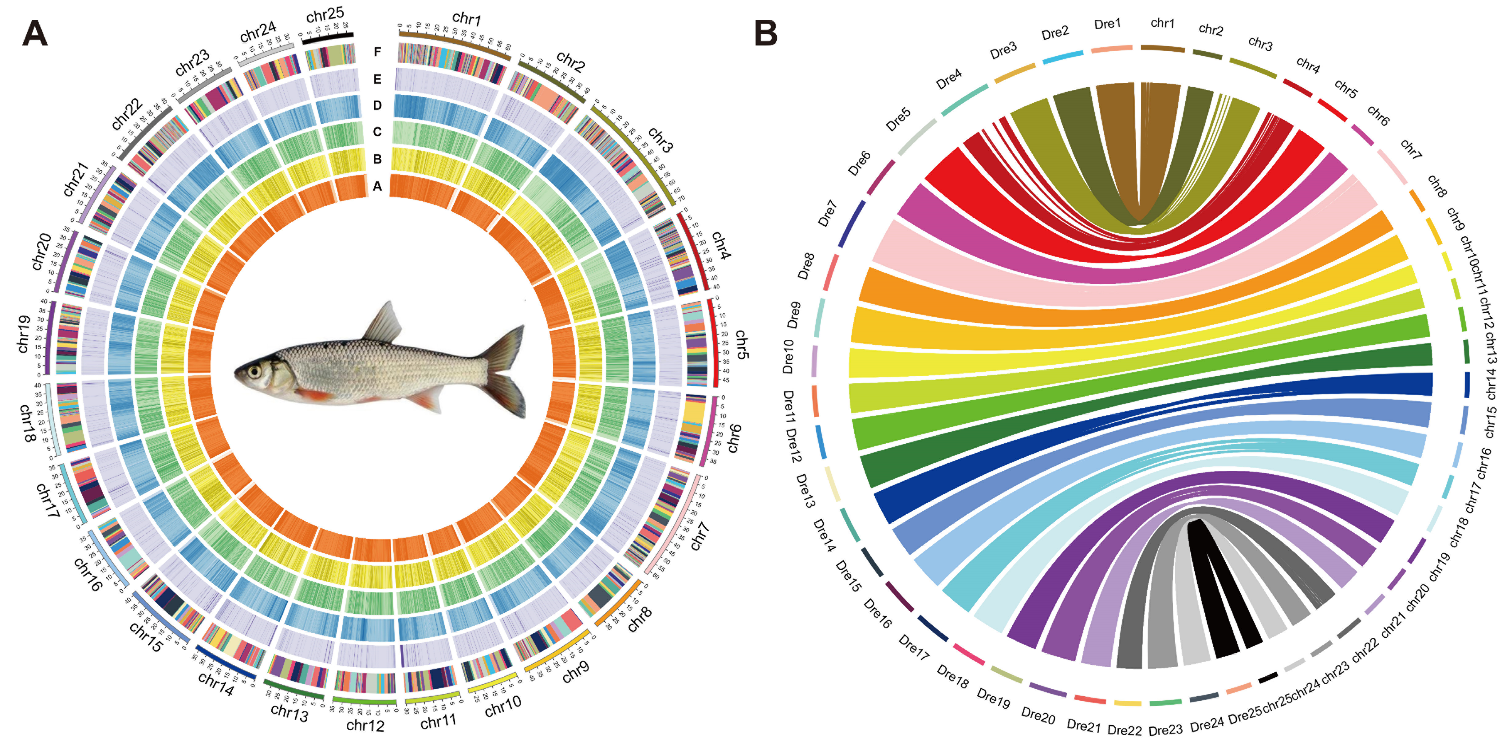

#### Fig S6. The genome landscape of Amur Ide and the genomic collinearity between Amur Ide and Zebrafish.

**(A) Circos plot of 25 chromosome-level scaffolds, representing annotation results of genes, ncRNAs and transposable elements on these scaffolds.** The tracks from inside to outside are gene abundance of the positive strand (red), gene abundance of the negative strand (yellow), TE abundance of positive-strand (green), TE abundance of the negative strand (blue), the ncRNA abundance of both strands (purple), and contigs which comprised the scaffolds (adjacent contigs on a scaffold are painted in different colors). **(B)** **Circos diagram between *L. waleckii* and *D. rerio*.** Each colored arc represents an 1 Kb fragment match between two species. We messed up the order of the Amur Ide chromosomes on the image for better illustrate our results.

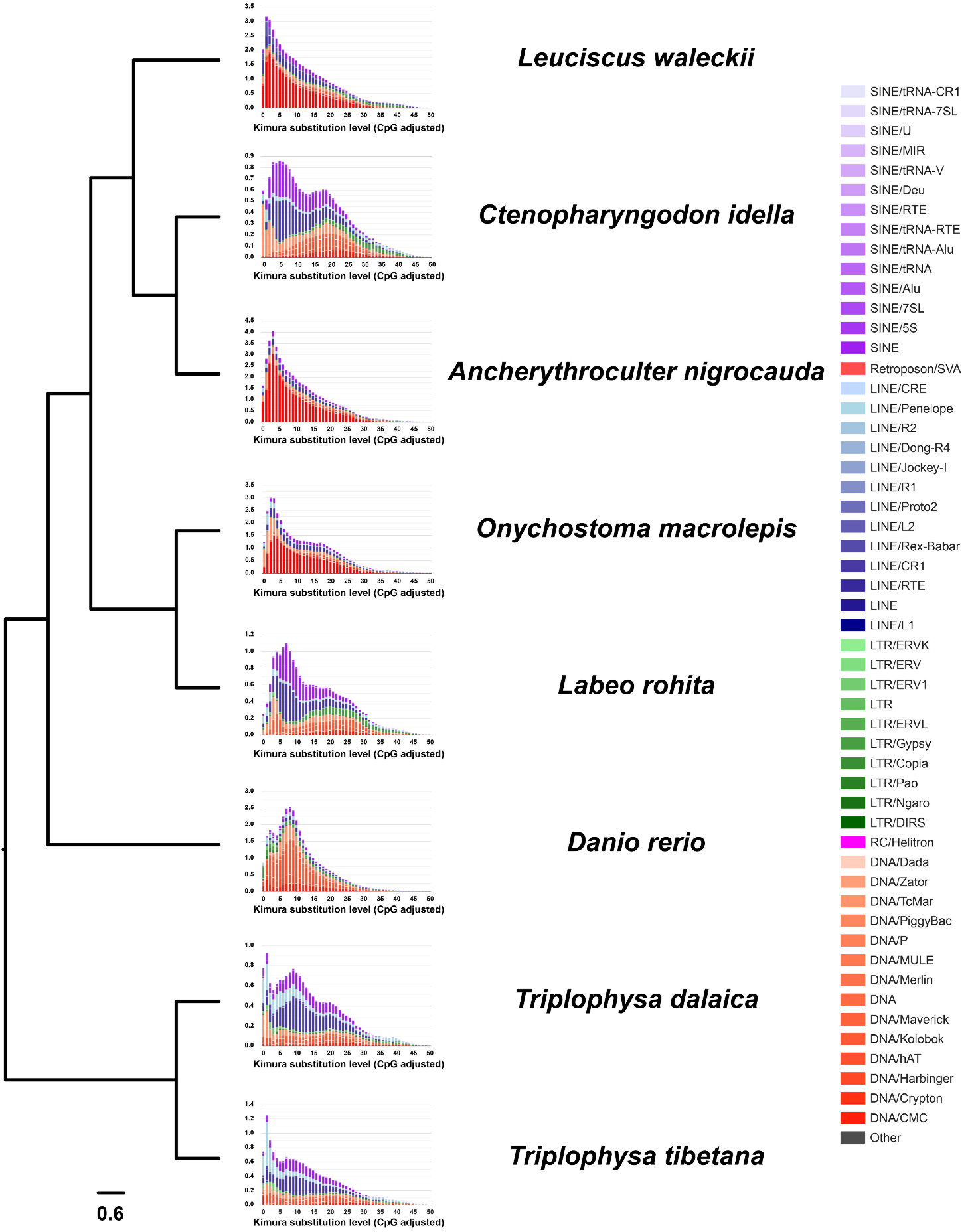

#### Fig S7. The divergence distribution of TEs in the Amur Ide and related species.

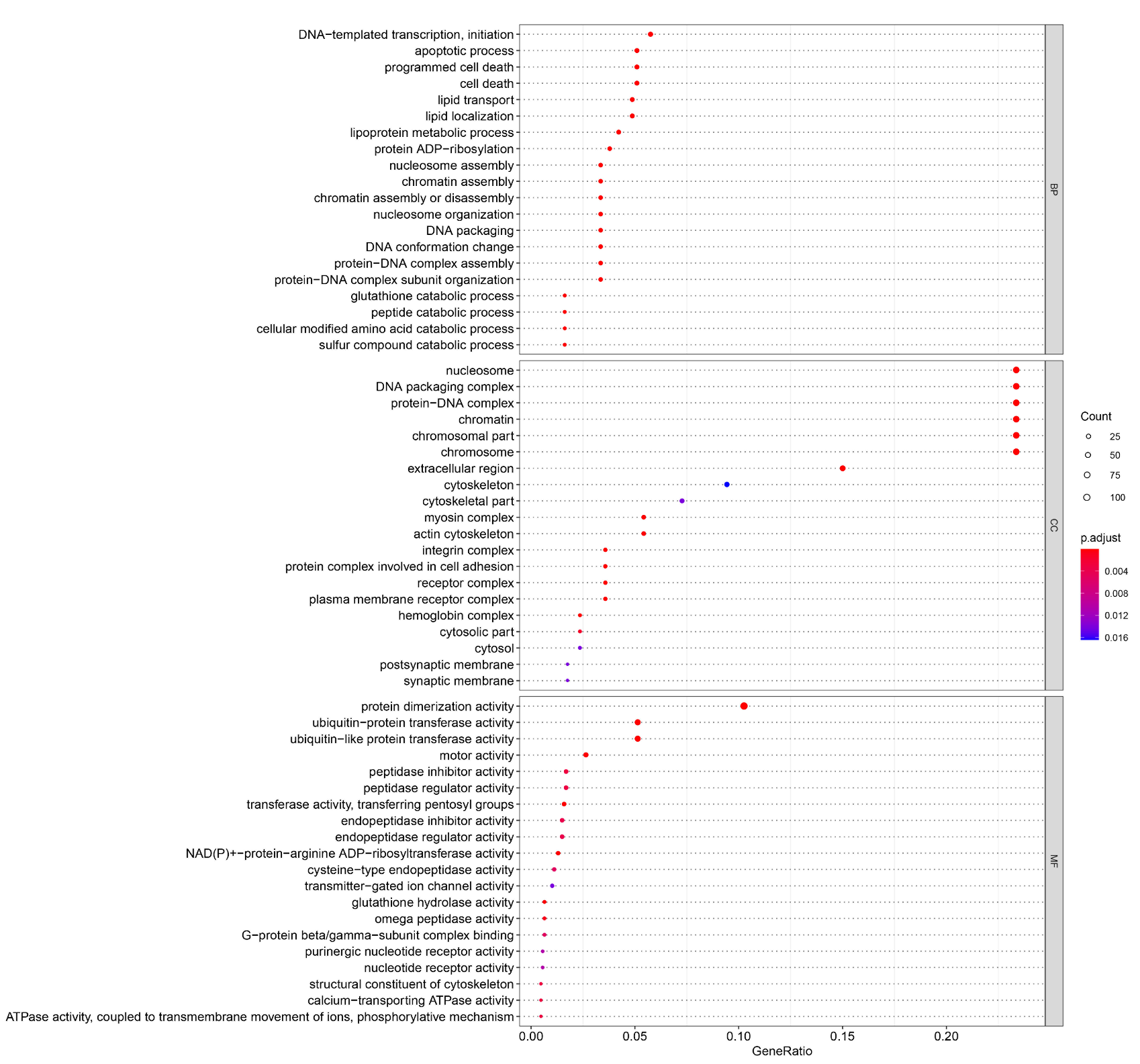

#### Fig S8. The bubble diagram of GO enrichment of expansion gene families in Amur Ide.

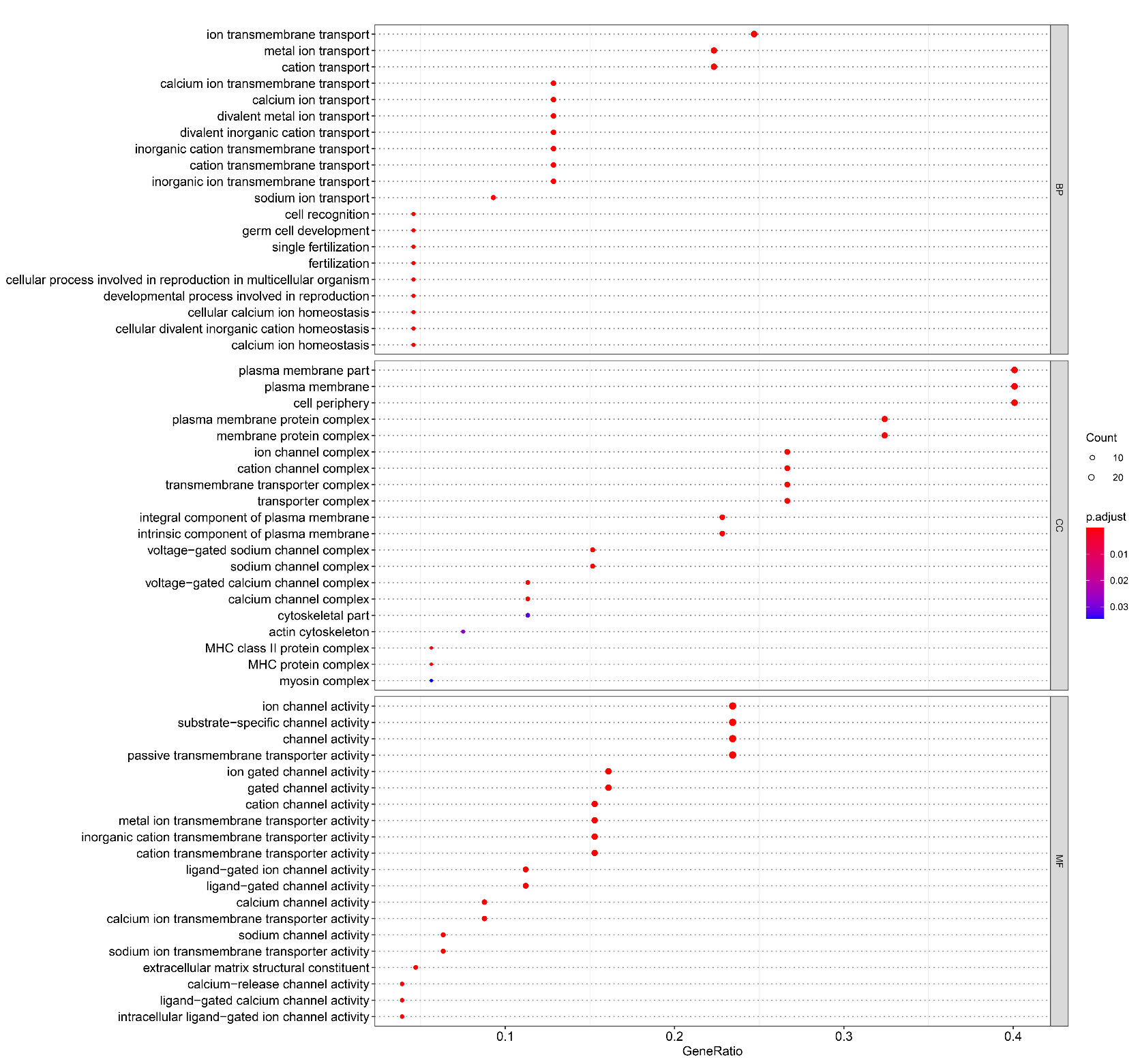

#### Fig S9. The bubble diagram of GO enrichment of contraction gene families in Amur Ide.

#### Fig S10. Distribution of monsoon in Northeast Asia and the location of Lake Dali Nur.

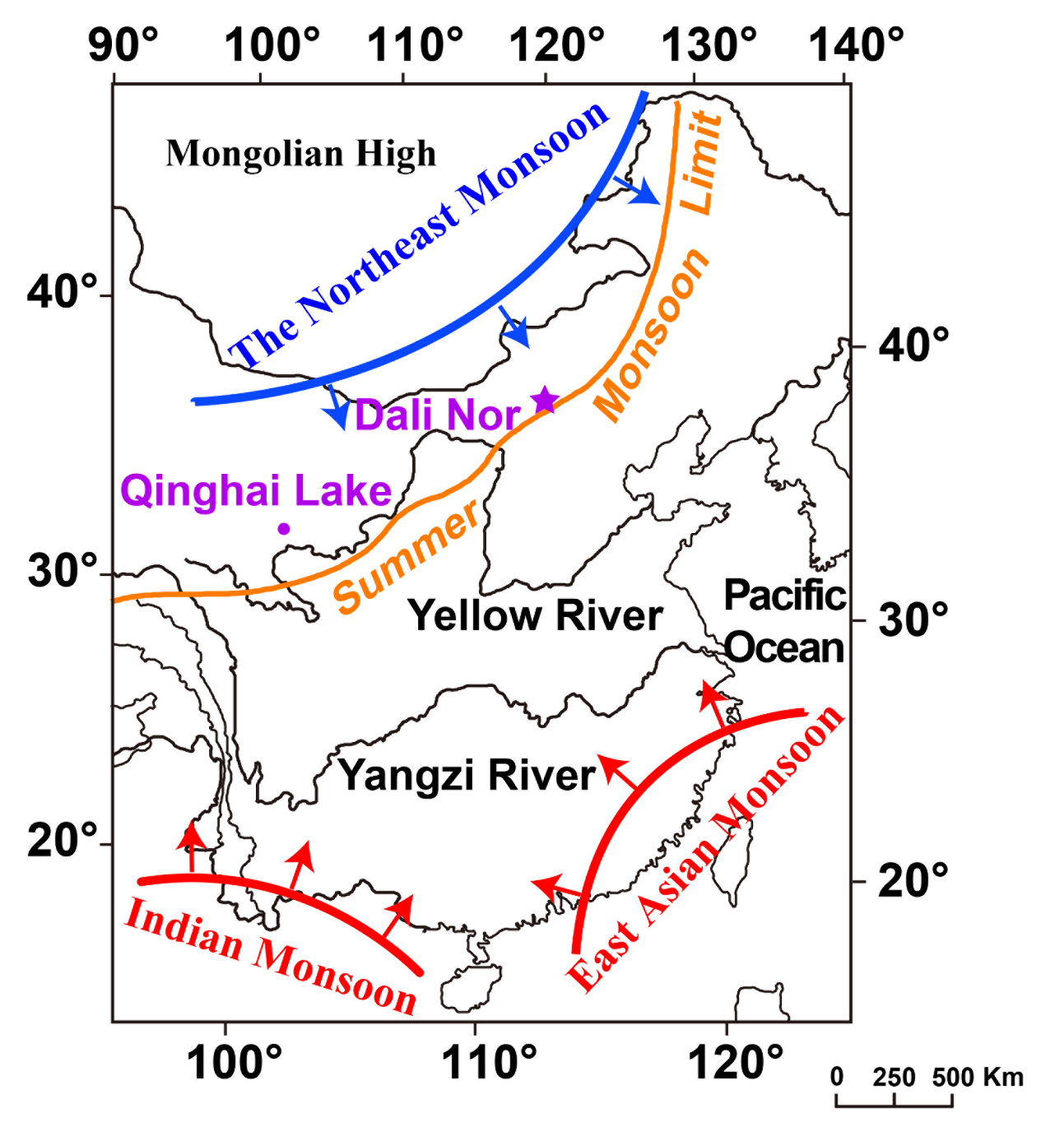

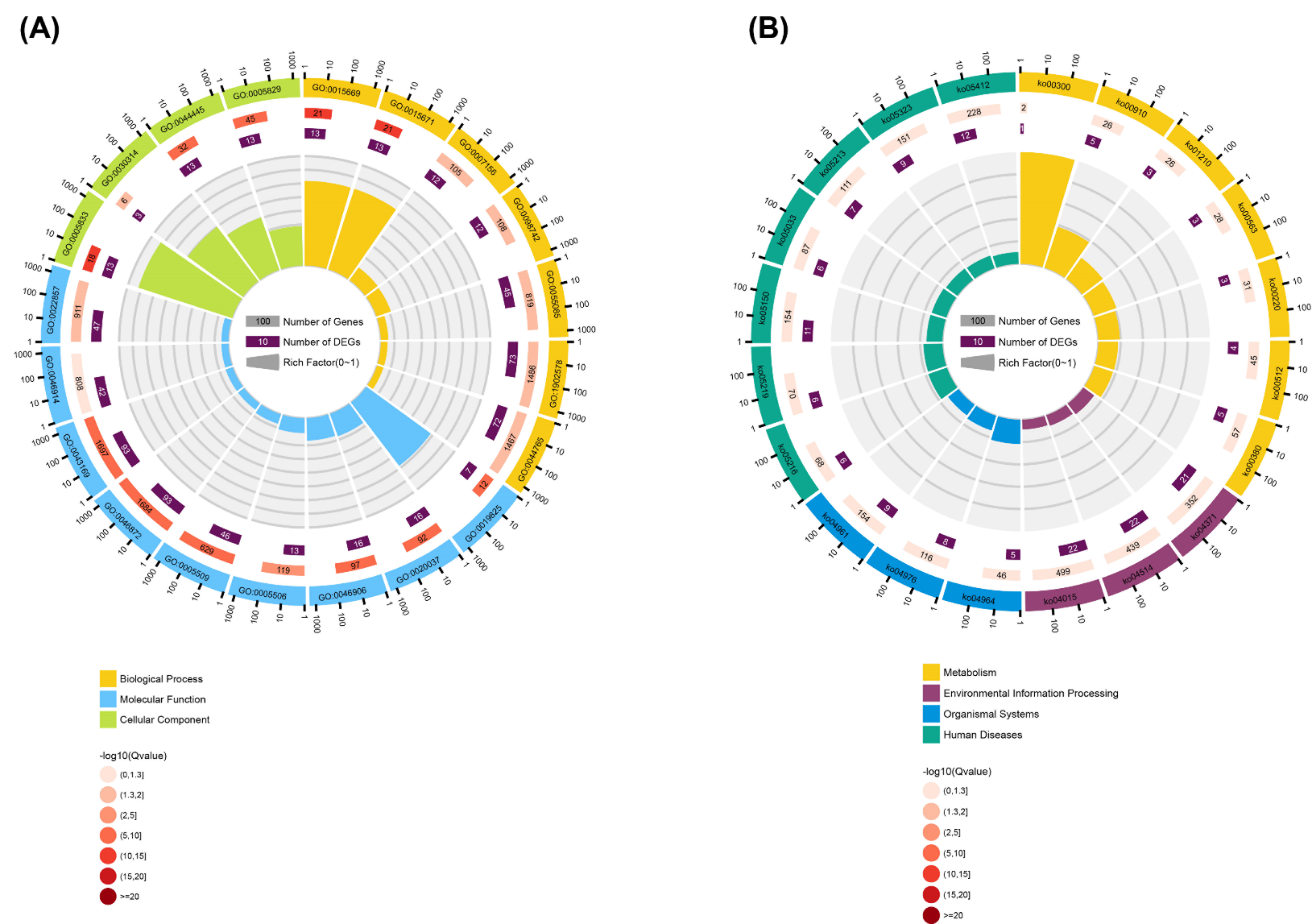

#### Fig S11. The enrichment of candidate selected genes.

**(A) The GO enrichment of candidate selected genes identified by Fst. (B) The GO enrichment of candidate selected genes is identified by π ratio.**

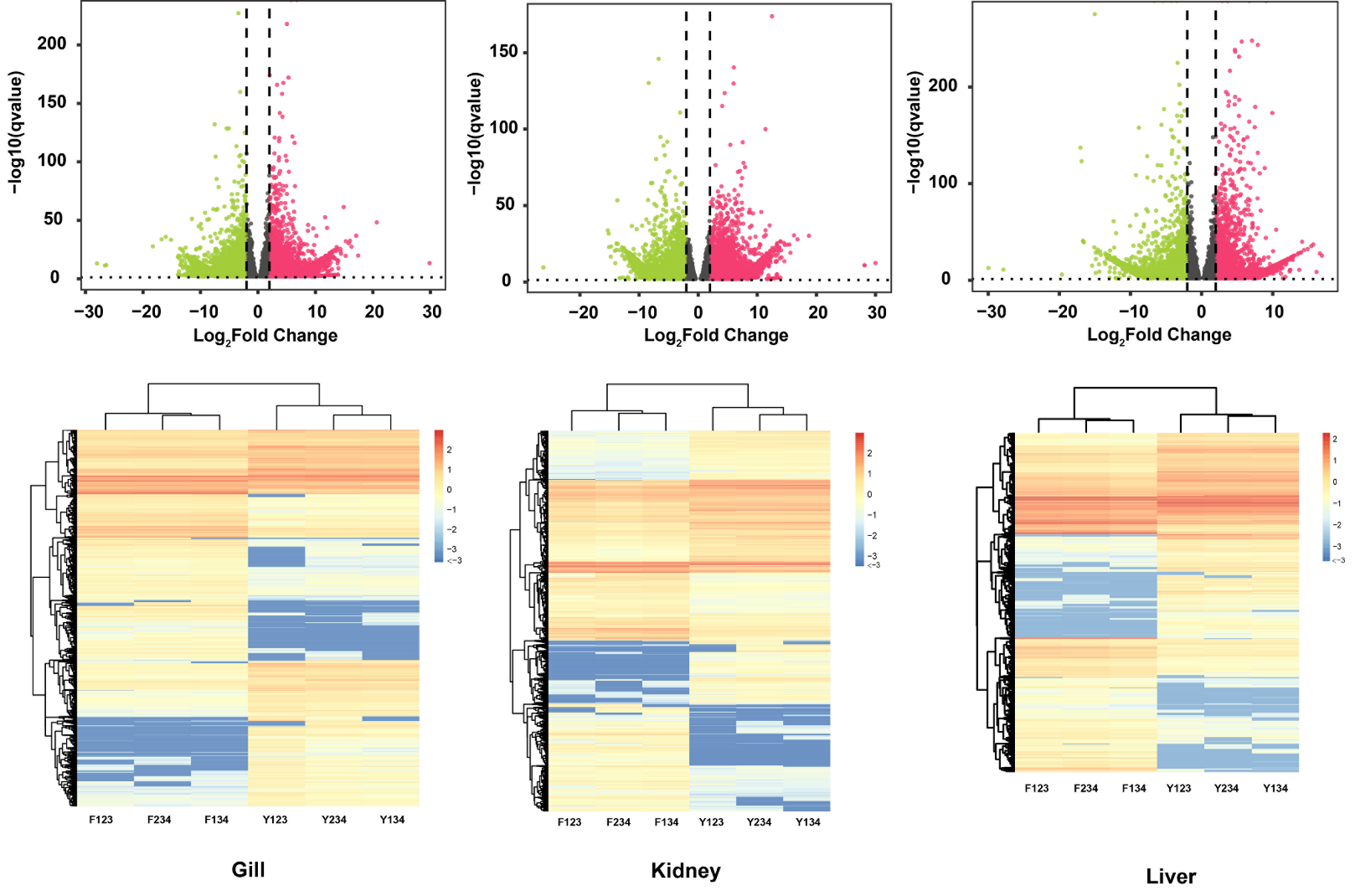

#### Fig S12. The volcanic map and the heatmap of differential expression analysis in the gill, liver and kidney of Amur Ide.

#### Fig S13. The phylogenetic tree of CA genes between Amur Ide and releted species.

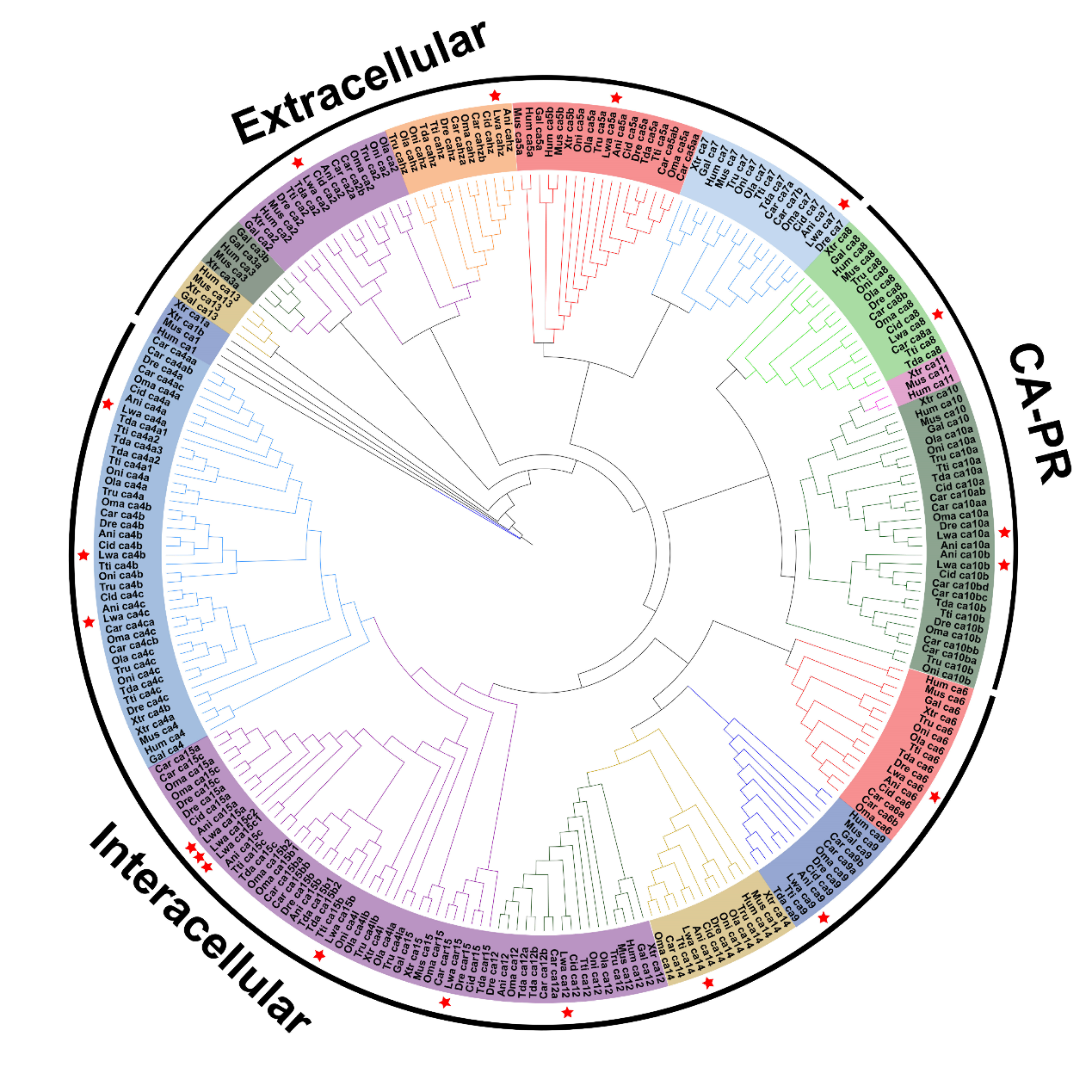
19 CA genes of *L. waleckii* were marked by the red star.

#### Fig S14. The motif of 19 CA genes in Amur Ide.

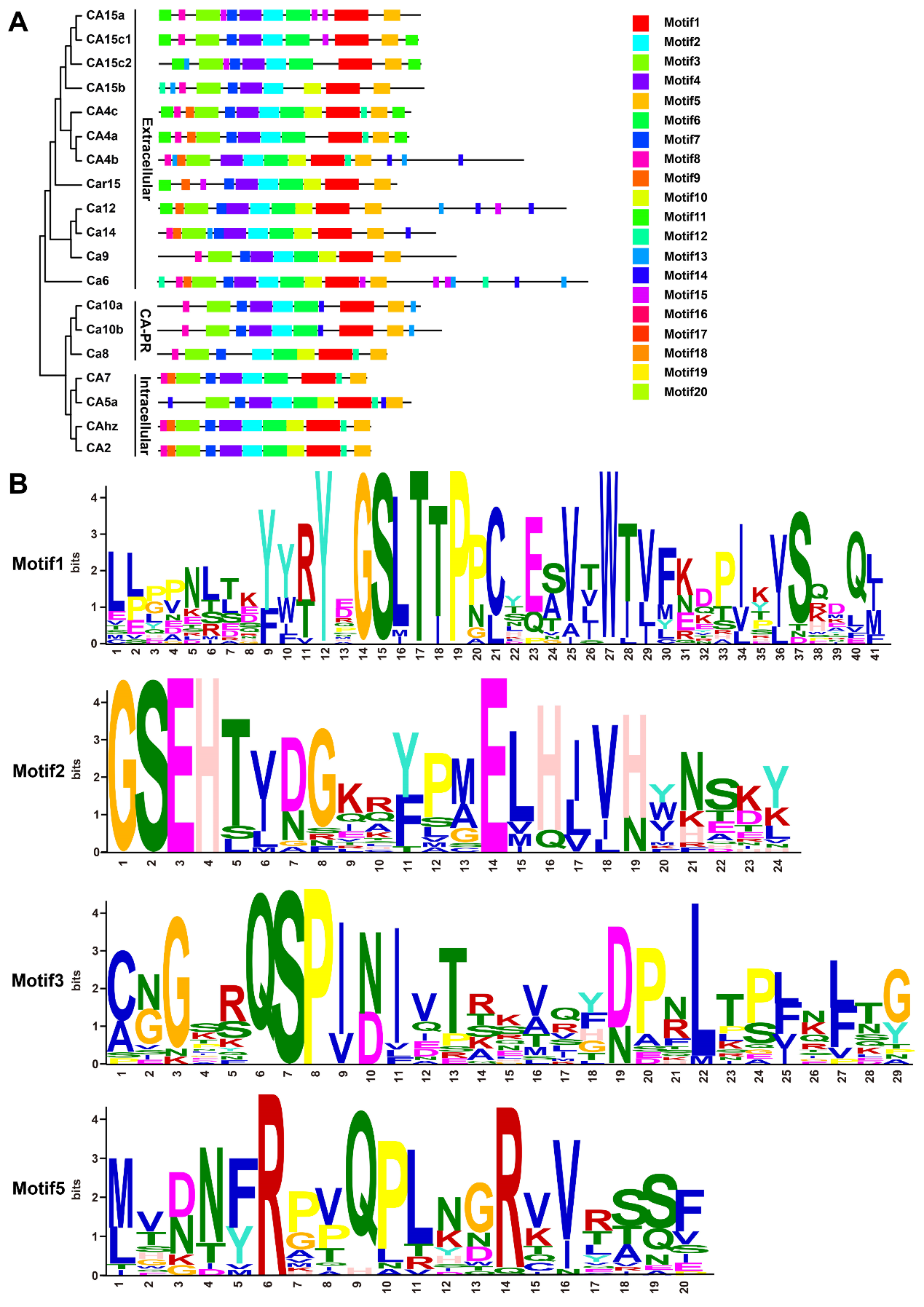

(A) The motif distributions of 19 CA genes in Amur Ide; (B) The shared 4 motifs of 19 CA genes in Amur Ide.

#### Fig S15. The phylogenetic tree of RH glycoproteins between Amur Ide and releted species.

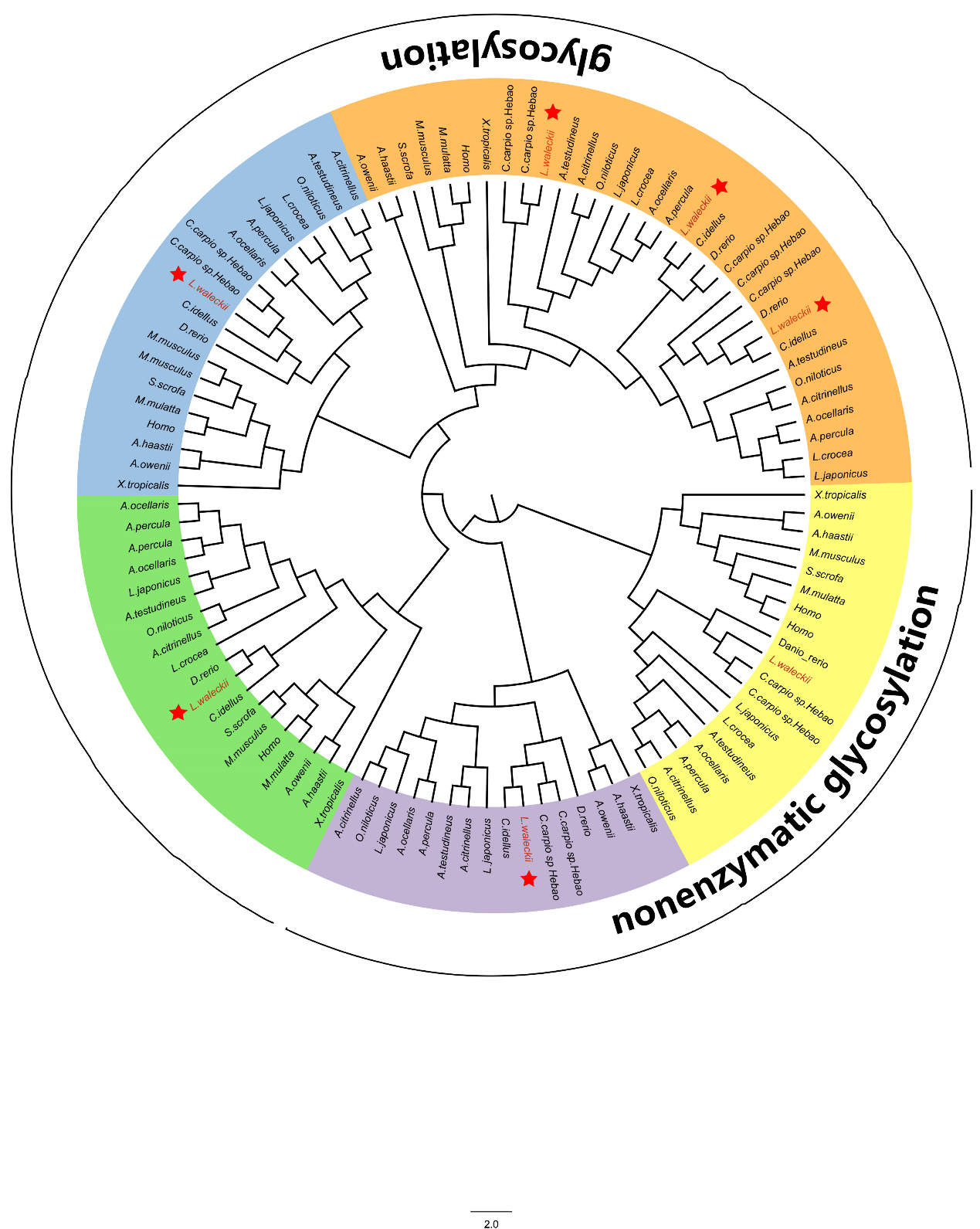

7 RH genes of *L. waleckii* were marked by the red star.

#### Fig S16. The motif of 7 RH genes in Amur Ide.

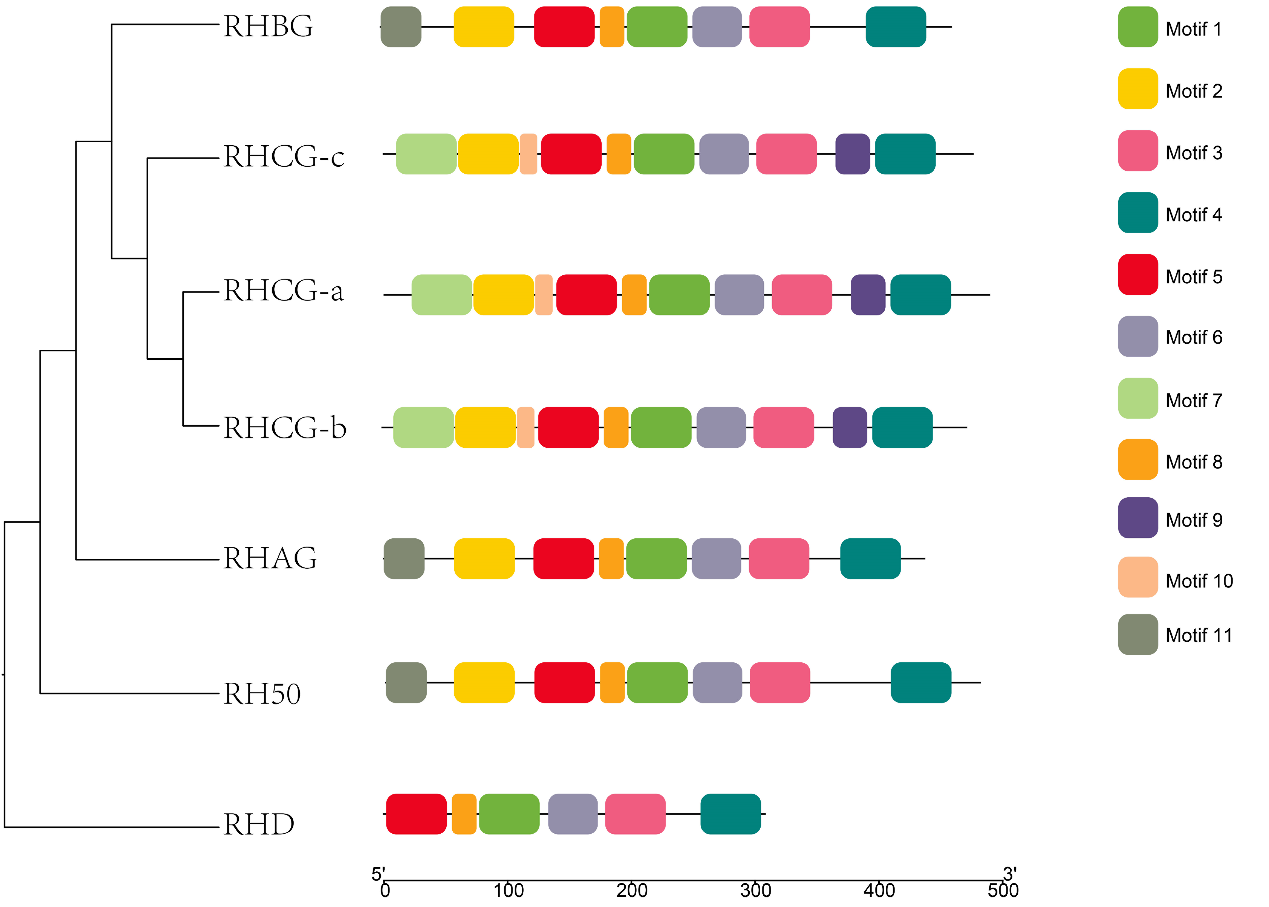

#### Fig S17. The 3D structure of RHCG-a of ALK and FW *L. waleckii* population.

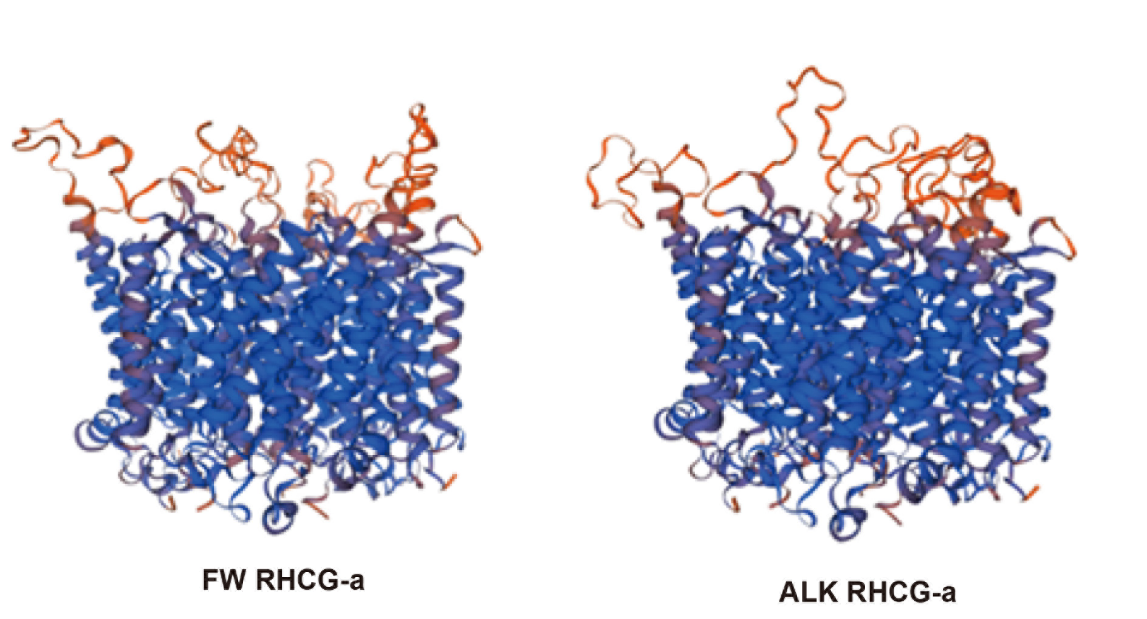

### Supplementary Table

**Table S1. The statistics of genome sequencing data.**

| Pair-end libraries | Insert size | Total data (G) | Read Length | Sequence coverage (X) |
| --- | --- | --- | --- | --- |
| Illumina reads | 350bp | 73 | 150 | 64.6 |
| PacBio reads | - | 62 | - | 54.87 |
| HiC | - | 102.93 | 150 | 92.35 |
| Total | - | 120.77 | - | 199.23 |

#### Table S2. The statistics of 17 K-mer analysis of *L. waleckii*.

| Depth | number of kmer | Genome_size(M) | Revised Genome_size(M) | Heterozygous_rate(%) | Repeat_rate(%) |
| --- | --- | --- | --- | --- | --- |
| 43 | 49238269810 | 1145.08 | 1125.03 | 0.56 | 57.61 |

#### Table S3. The statistics of PacBio Sequencing.

| Library | Read base (G) | Read number | Read length（Mean） | N50 |
| --- | --- | --- | --- | --- |
| PacBio | 73 | 2,030,960 | 35,943 | 62,499 |

#### Table S4. The statistics of assembly of Amur Ide genome.

|  | Contig Length (bp) | Contig number |
| --- | --- | --- |
| Total | 1,103,966,172 | 6,407 |
| Max | 12,092,634 | - |
| Length>=2000 |  | 6,406 |
| N50 | 1,515,867 | 180 |
| N60 | 1,017,170 | 270 |
| N70 | 681,883 | 405 |
| N80 | 336,993 | 630 |
| N90 | 95,719 | 1,246 |
| GC content | 0.3877 |  |

#### Table S5. The statistics of Repeat elements annotation of Amur Ide genome.

| Type | Repeat Size (bp) | % in genome |
| --- | --- | --- |
| Trf | 69,759,993 | 6.32 |
| Repeatmasker | 524,034,603 | 47.47 |
| Proteinmask | 82,731,199 | 7.49 |
| Total | 551,058,658 | 49.92 |

#### Table S6. Detailed classification of repeat sequences in Amur Ide genome.

(Included in a separated excel file)

#### Table S7. The statistics of gene structure annotation of Amur Ide genome.

(Included in a separated excel file)

#### Table S8. The statistics of gene function annotation of Amur Ide genome.

|  | Number | Percent (%) |
| --- | --- | --- |
| Total | 27633 | - |
| Swissprot | 22629 | 81.9 |
| Nr | 26579 | 96.2 |
| KEGG | 23536 | 85.2 |
| InterPro | 24392 | 88.3 |
| GO | 16660 | 60.3 |
| Pfam | 21205 | 76.7 |
| Annotated | 26613 | 96.3 |
| Unannotated | 1020 | 3.7 |

#### Table S9. The statistics of chromosome assembly of Amur Ide.

| chromosome | Length |
| --- | --- |
| chr3 | 71368983 |
| chr1 | 63702483 |
| chr7 | 62271240 |
| chr5 | 49313439 |
| chr4 | 45682451 |
| chr2 | 42701760 |
| chr9 | 41820828 |
| chr15 | 41008407 |
| chr19 | 40980234 |
| chr22 | 40598971 |
| chr18 | 40463193 |
| chr6 | 39639001 |
| chr14 | 37199493 |
| chr16 | 37024483 |
| chr21 | 36678393 |
| chr20 | 35751175 |
| chr12 | 35278759 |
| chr17 | 35180327 |
| chr8 | 34866635 |
| chr23 | 34564867 |
| chr13 | 34473392 |
| chr24 | 32144632 |
| chr11 | 30489377 |
| chr25 | 28728858 |
| chr10 | 28415676 |
| sum | 1020347057 |
| N50 | 39639001 |
| Total | 1105256174 |

#### Table S10. The BUSCO survey of Amur Ide.

| Species | BUSCO notation assessment results |
| --- | --- |
| *L.waleckii* | C:96.4%[S:93.6%,D:2.8%],F:2.4%,M:1.2%,n:4,584 |

#### Table S11. The mapping statistics of Illumina sequencing reads to reference Amur Ide genome.

|  |  | Percentage |
| --- | --- | --- |
| Reads | Mapping rate (%) | 99.05 |
| Genome | Average sequencing depth | 48.66 |
|  | Coverage (%) | 93.65 |
|  | Coverage at least 4X (%) | 92.19 |
|  | Coverage at least 10X (%) | 90.82 |

#### Table S12. The statistics of SNPs in the genome sequencing individual.

|  | Number | Percentage |
| --- | --- | --- |
| All SNP | 2,707,134 | 0.2756 |
| Heterozygosis SNP | 2,662,101 | 0.271 |
| Homology SNP | 45,033 | 0.0046 |

#### Table S13. The statistics of detailed classification of transposable element sequences between Amur Ide and related species*.*

(Included in a separated excel file)

#### Table S14. The statistics of the number and frequency of various amino acids used in protein-coding between Amur Ide and related species*.*

(Included in a separated excel file)

#### Table S15. The statistics of the number and frequency of various tRNA genes between Amur Ide and related species.

(Included in a separated excel file)

#### Table S16. The statistics of orthologues gene families between Amur Ide and related species.

(Included in a separated excel file)

#### Table S17. The statistics of expansion and contraction gene families between *L. waleckii* and related species.

| Species | Expansion | Contractions | Other | Total |
| --- | --- | --- | --- | --- |
| Tti | 1123 | 2705 | 21022 | 24850 |
| Tda | 1736 | 1372 | 21742 | 24850 |
| Dre | 4421 | 6930 | 13499 | 24850 |
| Lro | 3598 | 7706 | 13546 | 24850 |
| Oma | 1330 | 2732 | 20788 | 24850 |
| Lwa | 1751 | 5202 | 17897 | 24850 |
| Cid | 2733 | 6227 | 19384 | 24850 |
| Asg | 1395 | 8495 | 14960 | 24850 |
| Ani | 2270 | 3622 | 18958 | 24850 |
| Ola | 1965 | 10237 | 12648 | 24850 |

#### Table S18. The GO enrichment of expansion gene families in Amur Ide*.*

(Included in a separated excel file)

#### Table S19. The GO enrichment of contraction gene families in Amur Ide.

(Included in a separated excel file)

#### Table S20. The gene numbers of GGT gene family between Amur Ide and related species.

|  | Agr | Ani | Cid | Lwa | Oma | Dre | Tda | Tti | Ola |
| --- | --- | --- | --- | --- | --- | --- | --- | --- | --- |
| *ggt1* | 2 | 1 | 2 | 2 | 1 | 2 | 2 | 2 | 0 |
| *ggt1l* | 1 | 1 | 4 | 8 | 2 | 2 | 2 | 1 | 0 |
| *ggt5* | 2 | 1 | 2 | 2 | 1 | 2 | 2 | 2 | 0 |
| *ggt6* | 1 | 1 | 1 | 1 | 1 | 0 | 1 | 1 | 0 |
| *ggt7* | 0 | 1 | 1 | 1 | 1 | 1 | 1 | 1 | 3 |
| sum | 6 | 5 | 10 | 14 | 6 | 7 | 8 | 7 | 3 |

#### Table S21. The rapid evolution genes and function annotation in Amur Ide.

(Included in a separated excel file)

#### Table S22. The GO enrichment of rapid evolution genes in Amur Ide.

(Included in a separated excel file)

#### Table S23. The positively selected genes and function annotation in Amur Ide.

(Included in a separated excel file)

#### Table S24. The GO enrichment of positively selected genes in Amur Ide.

(Included in a separated excel file)

#### Table S25. The statistics of ReviGO analysis of rapid evolution genes in Amur Ide.

(Included in a separated excel file)

#### Table S26. The statistics of ReviGO analysis of positively selected genes in Amur Ide.

(Included in a separated excel file)

#### Table S27. The statistics of Genome Resequencing between ALK and FW Amur Ide population.

(Included in a separated excel file)

#### Table S28. The statistics of SNPs between 4 Amur Ide population.

| Category | DL | WS | YD | HL | ALL |
| --- | --- | --- | --- | --- | --- |
| SNPs | 3513387 | 3922726 | 3122707 | 2855556 | 6206224 |
| Intergenic SNPs | 2295142 | 2539536 | 2013242 | 1872427 | 4033100 |
| SNPs in Gene | 1218245 | 1383190 | 1109465 | 983129 | 2173124 |
| Intronic SNPs | 1061562 | 1220455 | 977574 | 869144 | 1929389 |
| Exonic SNPs | 156683 | 162735 | 131891 | 113985 | 243735 |
| SNPs in 3’-UTR | 24086 | 27000 | 21665 | 16668 | 41816 |
| SNPs in 5’-UTR | 3090 | 2830 | 2285 | 1493 | 5756 |
| SNPs in CDS | 129507 | 132905 | 107941 | 95824 | 196163 |
| No. INDELs | 110316 | 274921 | 368693 | 416994 | 845427 |

#### Table S29. The statistics of Fst and π ratio scanning in 25 chromosomes between ALK and FW Amur Ide population.

| Chromosome | median of π | | median of ρ | |
| --- | --- | --- | --- | --- |
|  | ALK | FW | ALK | FW |
| 1 | 0.000981186 | 0.00102222 | 109.5369838 | 142.8797446 |
| 2 | 0.00113694 | 0.00124035 | 110.456985 | 163.4349425 |
| 3 | 0.00106511 | 0.001086 | 130.6563018 | 147.8673014 |
| 4 | 0.00107276 | 0.0012121 | 110.7087671 | 158.3597406 |
| 5 | 0.00112776 | 0.00118218 | 109.4068606 | 157.323508 |
| 6 | 0.00101668 | 0.00112334 | 92.76513626 | 160.3161602 |
| 7 | 0.00105044 | 0.00109264 | 109.8517279 | 153.8119188 |
| 8 | 0.00119754 | 0.0012819 | 113.9412327 | 169.3761975 |
| 9 | 0.00110195 | 0.00118365 | 102.2559117 | 158.5018678 |
| 10 | 0.00120777 | 0.00136961 | 105.4763692 | 177.1545242 |
| 11 | 0.00104441 | 0.00120533 | 90.83679528 | 172.8768996 |
| 12 | 0.000965553 | 0.00112858 | 82.60421771 | 167.1634125 |
| 13 | 0.0012115 | 0.00125448 | 105.1378618 | 173.3711374 |
| 14 | 0.0011941 | 0.00121113 | 121.3783641 | 166.766422 |
| 15 | 0.00110581 | 0.0011551 | 116.7737874 | 154.4111877 |
| 16 | 0.00112454 | 0.00119826 | 103.3940643 | 166.9325362 |
| 17 | 0.00115974 | 0.00122642 | 101.1232752 | 168.8686508 |
| 18 | 0.00110132 | 0.00119132 | 101.1886091 | 160.6208974 |
| 19 | 0.00114627 | 0.00114823 | 119.879374 | 161.8801581 |
| 20 | 0.00100791 | 0.00116224 | 94.20244469 | 166.4233426 |
| 21 | 0.00110343 | 0.0011886 | 102.7857523 | 164.0646555 |
| 22 | 0.00108238 | 0.00113934 | 133.3337419 | 157.3361366 |
| 23 | 0.000993889 | 0.0011473 | 84.33045281 | 164.3682196 |
| 24 | 0.00104489 | 0.00117884 | 99.18769828 | 164.6084054 |
| 25 | 0.00119587 | 0.00124743 | 117.9801652 | 167.1187866 |
| Average | 0.00120804 | 0.00128831 | 121.513 | 150.838 |
| P-value of T-test | 1.10E-04 | | 1.01E-23 | |

#### Table S30. The candidate selected regions that were identified by Fst in ALK Amur Ide population.

(Included in a separated excel file)

#### Table S31. The candidate genes that were identified by Fst in ALK Amur Ide population.

(Included in a separated excel file)

#### Table S32. The candidate selected regions that identified by π ratio in ALK Amur Ide population.

(Included in a separated excel file)

#### Table S33. The candidate genes that were identified by π ratio in ALK Amur Ide population.

(Included in a separated excel file)

#### Table S34. The GO enrichment of candidate selected genes in ALK Amur Ide population.

(Included in a separated excel file)

#### Table S35. The KEGG enrichment of candidate selected genes in ALK Amur Ide population.

(Included in a separated excel file)

#### Table S36. The statistics of RNA sequencing between ALK and FW Amur Ide population.

| sample | Tissue | raw_reads | clean_reads | clean_bases | error_rate | Q20 | Q30 | GC_pct |
| --- | --- | --- | --- | --- | --- | --- | --- | --- |
| Y123 | Gill | 23839355 | 23341780 | 7G | 0.03 | 97.84 | 93.83 | 40.86 |
| Y234 | Gill | 26461363 | 25944003 | 7.78G | 0.03 | 97.81 | 93.81 | 44.38 |
| Y134 | Gill | 24263567 | 23811428 | 7.14G | 0.03 | 97.88 | 93.99 | 43.75 |
| Y123 | Kidney | 23805989 | 23450813 | 7.04G | 0.02 | 98.06 | 94.39 | 45.29 |
| Y234 | Kidney | 22718146 | 22295627 | 6.69G | 0.03 | 97.96 | 94.11 | 45.58 |
| Y134 | Kidney | 23641964 | 23269140 | 6.98G | 0.02 | 98.23 | 94.69 | 45.17 |
| Y123 | Liver | 23113754 | 22706268 | 6.81G | 0.02 | 98.25 | 94.81 | 46.7 |
| Y234 | Liver | 24854195 | 24405169 | 7.32G | 0.02 | 98.49 | 95.28 | 46.9 |
| Y134 | Liver | 25939636 | 25413351 | 7.62G | 0.02 | 98.41 | 95.17 | 46.86 |
| F123 | Gill | 23851172 | 23304759 | 6.99G | 0.02 | 98.3 | 94.94 | 45.7 |
| F234 | Gill | 23483358 | 22992644 | 6.9G | 0.02 | 98.13 | 94.54 | 45.82 |
| F134 | Gill | 27551393 | 27010135 | 8.1G | 0.02 | 98.15 | 94.53 | 46.17 |
| F123 | Kidney | 23977208 | 23258985 | 6.98G | 0.02 | 98.27 | 95.03 | 46.75 |
| F234 | Kidney | 23263563 | 22797508 | 6.84G | 0.03 | 97.95 | 94.24 | 46.49 |
| F134 | Kidney | 23284498 | 22866900 | 6.86G | 0.02 | 98.35 | 95.07 | 46.55 |
| F123 | Liver | 23754957 | 23306632 | 6.99G | 0.03 | 97.69 | 93.37 | 46.63 |
| F234 | Liver | 23416841 | 23029748 | 6.91G | 0.02 | 98.49 | 95.22 | 46.79 |
| F134 | Liver | 27891699 | 27203087 | 8.16G | 0.02 | 98.49 | 95.25 | 47.14 |
| sum | - | 439112658 | 430407977 | 129.11G | - | - | - | - |

#### Table S37. The statistics of gene expression in gill between ALK and FW Amur Ide population.

(Included in a separated excel file)

#### Table S38. The up-regulated gene list of ALK Amur Ide population in gill.

(Included in a separated excel file)

#### Table S39. The down-regulated gene list of ALK Amur Ide population in gill.

(Included in a separated excel file)

#### Table S40. The statistics of gene expression in kidneys between ALK and FW Amur Ide population.

(Included in a separated excel file)

#### Table S41. The up-regulated gene list of ALK Amur Ide population in the kidney.

(Included in a separated excel file)

#### Table S42. The down-regulated gene list of ALK Amur Ide population in the kidney.

(Included in a separated excel file)

#### Table S43. The statistics of gene expression in liver between ALK and FW Amur Ide population.

(Included in a separated excel file)

#### Table S44. The up-regulated gene list of ALK Amur Ide population in the liver.

(Included in a separated excel file)

#### Table S45. The down-regulated gene list of ALK Amur Ide population in the liver.

(Included in a separated excel file)

#### Table S46. The copy numbers of CA genes between Amur Ide and related species.

#### Table S47. The accession number of CA genes in Cyprinidae fish.

(Included in a separated excel file)

#### Table S48. The statistics of 19 CA genes in ALK Amur Ide population.

(Included in a separated excel file)

#### Table S49. The statistics of highly differentiated sites in CA genes between ALK and FW Amur Ide population.

(Included in a separated excel file)

#### Table S50. The accession number of RH genes in Cyprinidae fish.

(Included in a separated excel file)

#### Table S51. The statistics of 7 RH genes in ALK Amur Ide population.

(Included in a separated excel file)

#### Table S52. The statistics of highly differentiated sites in RH genes between ALK and FW Amur Ide population.

(Included in a separated excel file)
