## Supplementary material for "Genomic insights into the adaptive and convergent evolution of *Leuciscus waleckii* inhabiting extremely alkaline environments": Supplymentary File3

>Ani|CA14

MDLVWITVSLSLFLKCVECTGTGKTLIIYVGAVGQPEWAEFFPDCGGSSQSPINLDKSQTRHDPTLIPVQPLGYNQPGNRPFTLSNNGHTVQMTLPHWMGLAGLPWHYSAVQLHLHWGNGVGVATGSEHTINGQSTSAEVSQLHIVHYNAEVYANMSQAKTQENGLAVLGILIEVGEEINQAYGSILNYLGRIRYAGQKVAIPAFDIQSLLPDNLSQYFRYNGSLTTPPCHQSVLWTIFNERVKISHSQVQRMKLETVLYSSKAEEAEPMVLQDNYRATQPLNHRTVLSSFIPVSGEITAIVIGSLCGCIGLAVIIYFIVKTIRLTSSWDVLPSIRPAPQPR

>Cid|car15

MMLLLMLMLSVVLLARSDDDFCYDEEHCDPYAWGDNYPSCHPLLDSHHSPINLDHQLMKNDSLDSLNLEGFNLTHKGQWWLTNQGHSVVLEVGNGMQVSGGGLPGTYRTIQLHFHWGSVSSNGSEHTLDHLRFPMEMHIVNIKSTHPNLTSALEDPTGIAVLGIFVDVTYLHNENFQSISSALPSVAYKGQTKSIRPFPLVNLLPQNNLTQYYRYHGSLTTPPCSQVVLWTVYEVPVYISWTQFEQFVSGIYSTEEEADNHVLLHDNYRHIHPTYGRSVYASKDAKLLTSGVSSLALFNLLNVLLLLIFKHLAFCI

>Cid|ca6

MTNWGTFISLPATKLWSLQKGELDQKHWAEKFHECGGNQQSPIDIQRRKVRYNPRMLQLELTGYEDMRGSFLMKNNGHSVEIQLPHTMMITKGFPDHYSAVQMHLHWGGWDLEASGSEHTMDGIRYMAELHVVHYNSEKYSSFQEAKDKPDGLAVLAFFYEDGHFENTYYSDFLANLANIKYAGQSMNITNLNVRSMLPENLNHFFRYQGSLTTPPCHESILWTVFDTPITLSHNQIRKLESTLMDHENKTLWNDYRMAQPLNDRVVESSFLPRLGKGGICRQEEIEAKLKRIESLIVSLDKKTVPGPQPLSPLVLYFAEKNLETYALVNLSHPMNLQSFTACMNVRIPPIQDLTVLSYSTSRDNELMITLGSEVGLWIGDEFVNLPFERQSHDWTNYCFTWASHTGGAELWVNGLVGDERYIRAGYTIPAGGKLILGKDQDGFLGISDTDAFVGHMTDVNVWDYVLTAAEIGGQMLCDNGKMKGNVLNWGTTQLSLYGGVQLQAEQVCH

>Tda|CA12a

MKTNTRHLFKMTLTLVFIILVGCPFACRGAKWTYNGPDGEHHWPRNYPFCGGAFQSPIDFQTQLLRYDPNLPPIQVQNYNLSAHEQLTLSNNGHSVQLSLPPHMYISSLPHRYSAAQLHFHWGSSNLLTGSEHTVNGKQFAAEMHVVHFNSEKYPNISMAVDKHDGLAVLGVFIEIGEYNPAFNKFFKYINGIRYKGQRIQVPSFDIRQLLPAALNEYYRYDGSLTTPPCYPSVLWTVFKKPVTVSHKQFLALATALYATYSQDSAPMPLHGNYRKPQLTDSRVVLVSFNDGALLTLTITSLVGAVLIIIIAWCLLKMRCKCDGQKETNSEYKPAQKKKAPPKYKLHI

>Tda|CA4c

MLITSAHTHAGCPSYEGIDSPVTRSGKHPPIYETGVTFILEDCCNLLSLSYSTEPLTAALTLGAISERRVADEKLTFDPGQNSIWGDEAEMYLSVTAHLISTPKCTIARDTAPRHTSPRTALTSDQEGNPVRWLNDYQEILNQSKHACIQPQNKQNCEWCYKSQFSCNDTCKEPDHWHKIFPKCGGQSQSPINIVTRKVQHNSNLTGFIFEGHEDCVNITVENHGHSAHFTLPPSVRLRGGGLPTTYKAVQFHLHWGEEQEAGSEHSVDGERYPMELHIVHIKEQYSSLQEAESDTTGVALLAFFFEVTLKPNKHFDRVIEALGRVRYHGNTSAISGFRLADILLPASRLSYYRYSGSMTTPGCDQIVVWTVFHQTLPISQKQLALVTEQLLFRTEKPMTGIFRPMQNLNGRIVFKSVKSDSVCVLPGLISLFLCLFCALGQQRCFG

>Tda|CA12b

MKTNTRHLFKMTLTLVFIILVGCPFACRGAKWTYNGPDGEHHWPRNYPFCGGAFQSPIDFQTQLLRYDPNLPPIQVQNYNLSAHEQLTLSNNGHSVQLSLPPHMYISSLPHRYSAAQLHFHWGSSNLLTGSEHTVNGKQFAAEMHVVHFNSEKYPNISMAVDKHDGLAVLGVFIEIGEYNPAFNKFFKYINGIRYKGQRIQVPSFDIRQLLPAALNEYYRYDGSLTTPPCYPSVLWTVFKKPVTVSHKQFLALATALYATYSQDSAPMPLHGNYRKPQLTDSRVVLVSFNDGALLTLTITSLVGAVLIIIIAWCLLKMRCKCDGQKETNSEYKPAQKKKAPPKYKLHI

>Tda|CAhz

MWLDMILESVCSAPDSDSESTSVNAAVNVEHWNKSQKIMSHSWGYGAHNGPEKWEEDFPIANGPRQSPIDIVPNQAQHDPSLKLLKLKYDPATAKGILNNGHSFQVDFDDEDDSSTMTGGPITGTYRLRQFHFHWGDSDDRGSEHTIAGNMFPSELHLVHWNTKYPSFGEAASQPDGLGVVGVFVKIGSANPRLQKVIDAFDDIKSKGKQTTFSNFDPKTLLPASLDYWTYDGSLTTPPLLESVTWIVLKEPISASPGQIAKFRSLLFTAEGETPCCMVDNYRPPQPLKGRKVRASFK

>Tda|CA7

MTGHHWGYGEANGPAEWHKGYPLAQGSRQSPIDIVPGSAVYDTSLSPISLFYDNCTSINITNNGHSVVVEFVDIDDRSVIQGGPLDNMYRLKQFHFHWGGKGCGGSEHTVAGKIYPSELHLVHWNSVKYKSFGEAAAAPDGLAVLGIFLEIGIEHKGLHQITDAMYMVKFKGTVADFKGFNPKCLLPLSLDFWTYPGSLTTPPLFESVTWIVLKDPLFLSEKQMGKFRTLNFNGEEEESRRRMENNFRPPQPIKGRTVRASFR

>Tda|CA4a1

MNVLWEESSCSLTDILEQKRRSDSQRVTFGFIAGVTGQSVTMKSLISLFLLSFVLHLSSSADWCYRTQVSCDGHCKGPERWSEVKADCGKGRQSPINIVTKKTKLDERLTPFKFTRYQDAFDSTITNNGHSVQVNILNAPTVSGGNLGYTYKAVQLHLHWGTDGGPGSEHTVDGEQYPMELHIVHMKQRYNSVQDALKDPSGIAVLGFFYEESKTPNKHYDQFIHALKSIQNTNGNVTLRKISLNQFILSEENMTNYYRYEGSLTTPGCTEAVVWTVFENPIPLDKEQLRAFSSLKFHDGKPMVGTFRPVQSRNGRTVYRSSGPAVLACTVLLFVSITTTLSLSHIN

>Tda|CA4a2

MTDRRMLVILTSFFSHGFVCHRYCDLTGDTGQSVTMKSLISLFLLSFVLHVFSSEEWCYQTKGSCKGPENWKEINETCGGNRQSPVNIVTKATQLDKRLTSFNLTGYKTLFDSTVKKNSHTIEVTINTTATVSGGNLEDTYKPLQFHLHWGTNGVRGSEHTIDGEQYPMELHVVHIKQKYKDLAEAFKDPFGVAAFGFFLEESNSDNKNFDNLINVLGSIKSKSGDVSIKNISASQFILPEENMTSYYRYNGSLTVPNCYESVIWTVFEKTIPLSKKQLEFFSSLKLDDGTPMVGTFRPPQPRSGRTVYRSSSPAVLASVVLLFISITTTFSLSHLC

>Tda|CA4a3

MKSLISLFLLSFVLHVFSSAGPENWKEINNNCGGDRQSPVNIVTKTTQLDKRLTSFNLTGYKTLFDSTVKKNSHTVEVTLNPTATVSGGNLEDTYKPLQFHLHWGTNGVRGSDHTIDGEQYPMELHVVHIKQKYSNLTEAFQDQSGLAVFGFFLEESNSTNKNFDNLINVLRSIENKSGDVSIKNISASQFILPEENMTSYYRYNGSLTAPNCSESVIWTVFEKTIPLSKKQLEFFSSLKLDDGTPMVGTFRPPQLRSGRTVYRSSSPAVLASVESNSDNKNFDNLINVLRSIENKSGDVSIRNISASQFILPEENMTSYYRYNGSLTAPNCSESVIWTVFEKTIPLSKKQTGYSPLPWKVVMNSPVIRLELKHTSTLIV

>Tda|CA9

MLLEVLFILHVLKSQVLTVESSSSSSESDESSEPDEEHSKGSSHQQHWGYHDQDSWLSSYEDCGGKSQSPINVNTRQVTYDPRLPVIHLEGYDLTDRPPLSLLNNGHTLQLSLPNSMRIVGGFRQIYVAAQLHFHWGTTEVPGSEHTIDNIHFPAEIHVVHYNSKYPNLSEAASKADGLAVFGAFIGIGLYENDNYEKILSALTDISREESNTEVPAFNIRHLLPNSLERFYRYNGSLTTPPCFQTVSWTLFNDTIRVSRRQLAALEDTLKTEHNKLLSRNFRAPQLLHNRNVLTSFQTVAAAGSARGAHLSETEDANIKESSETQEAFSKGDVLAIVFGVLFAVTLLAFLVYAYQQRKKYSKYKRHSKQNVIYKPAVKEEA

>Tda|CA10a

PSFWGLVNSAWNLCSVGKRQSPVNIETSHMIFDPFLTPLRLNTGGRKVGGTMYNTGRHVSLRLDKEHLVNISGGPVTYSHRLEEIRLHFGSEDGQGSEHLLNGQAFSGEVQLIHYNHELYTNYTEAAKSPNGLVIVSIFMKISETSNSFLNRMLNRDTITRITYKNDAYLLSGLNIEEVYPETSSFITYEGSMTIPPCYETATWILMNKPIYVTKMQMHSMRLLSQNQPSQIFLSMSDNVRPVQPLNNRCIRTNINFSMQGKDCPNNRAQKLQYRVNEWLLK

>Tda|CA10b

MPHVWEFILILNINVIFLEAQPVSSKLHDGWWAYKDVVQGSFIPVPSFWGLVNTAWNLCAIGKRQSPINIETSRMIFDPFLGPLRLNVGQRKVSGTMYNTGRHVSLRPDKSHLVNISGGPLSYSYRLEEIRLHFGSEDSRGSEHLLNGQAFPGEVQLIHYNQDLYLNYSDAARGPNGIAVVSIFMKISEPTNSFLNRMLSRETVTRITYKHDAYLLMGLDIEELYPETSRFITYEGSITIPPCLETATWILMNRPIYISQIEMQSLRLLSLNQPSQIFLSMGDNMRPTQSLHQRCIRTNINFSQRRDCPNNRVMRPQYRVNEWLLK

>Tda|CA6

MELFQVLLYATFVNFASAGIDGAKWTYLDGELDQKHWAEKFPDCGKNQQSPIDIQRRKVRYNPRMLKLELSGYDEMHGQFVMTNNGHSVEIELPHTMMITKGFPGRYTAVQMHLHWGGWDMEASGSEHTVDGIRYMAELHIVHYNSEKYSSFKEAKDKPDGLAVLAFFYEDGHFENTYYSDFIANLAKVKYADQSMNISNLNVRSMLPENLNHFFRYQGSLTTPPCHESILWTVFDTPITLSHNQIRKLESTLMDHENKTLLNDYRMAQPLNDRVVESSFLPRLGKGGLCRQEEIESKLKKIENLIVSLDKKMLPGPPFPDRDLQTLFPLVLYFPEKNLETYALVNLTQPMTLQSFTVCMNVRTPPRNDLTVLSYSTSRNDNELMISLGSEVGLWIGNEFVNLPFDRQSHDWTHYCFTWA

SHTGGAELWVNGLVGEERYIKTRYTIPSGGTLILGKDQDGFLGISATDSFVGYMTDVNVWDHVLTVEEIKEQMLCSNGKVKGNVLSWGITQLSLYGGVQLSAEQVCH

>Tda|CA15b1

MKVQGGDLPGLFISTQIHFHWGKGSAMPGSEHAVDGKRYPMELHIVNKAVDDPNLLAALGFFIEATNDTGKPKSWKILTSYLAEIVNAGDKVCITKYISMDDLLPGVDRTKYYRYFGSLTTPNCIEGVVWTIFKDPIKVSRDLIDLFSTTVHVNKASSSPLMTNTFRGIQAINRRIVKSQVTGSILSSETNPCF

>Tda|CA15c

MERGSGGTFYQWSRGVQIRANLDLLIDWAHGAGLNDLAHSYLVKLSSAVNLMAKPKENLLQMSWSALRIEYNALNPAQLHHILREYNSQRSCPAAWTPSSDETTTALRTIDILEGFDNHPPLILPPDGFVFDLKRPISETGLVKQLSGFQRLIQKLPDSESVPDEPSQPASALPMKAKLTRMEVHPTAHLESGHTEIVFPHAEMNDWSGREAHLTKKLQNMELQKKGSHSRLEHSALDETCLLTPPNTPLNLEQMDSETDQQEEISYHQHFKYTCANKQEEGESEEEDVFVLELERGTCGLGLVVVDGQETQFKGNGMYIKSVMPDSPAALSQRLKAGDRILAVNGLSLVGVDYQTGRELIQASGDKPRLLESFITCKMIILLLCVVLLCPTVYSSDASVEWCYHEPRCDSTTWSKNVHD

YCSGTRQSPINIVTSNIMANSKLTPFNFTGFDDNSTFFSIGNTGDSVVVSLDGDKMTVEGGDLPSKYTSVQFHLHWGNGSSVNGSEHTVDGKRYPMELHIVNVHSKYNGNAKAAIDAKDPNGLAVLGFFVEGSSVTSKSKGWEILTSFLPNITNSSVTTTDIINTTTMNSLLEGVDRTKYYRYQGSLTTPRCNEIVIWTVFKDPIKVNHDLIQLFSTTVQFKGTSTKMANNFRGVQNMNGRIVTSQVASAASSTVIALTFYILAILPFICWL

>Tda|CA15b2

MKVQGGDLPGLLISTQIHFHWGKGSAMPGSEHAVDGKRYPMEIHIVNKAVDVPNLLAALGFFIEATNDTGKPKSWKILTSYLAEIANAGDKVCITKYISMDDLLPGVDRTKYYRYFGSLTTPNCIEGVVLTIFKDPIKVSRDLYNCPCQQGVQLTPNDQHFQGDPGHQSQNCQITKVHHQSSPHPHRLSANSDKQRSLFYQEKEKCRIIQSVLNVMGSFLGKTDNFR

>Tda|Car15

MKSMMLLLVMMLSGVLLAHSDDFCYGDERCDPYAWGDNYPSCHPLLESHHSPINLNHQLTKNGSLDSLNLEGFNLTPKGQWRLVNQGYSIVLEVGDGMQVSGGGLPGIYRTVQLHFHWGSVFTSGSEHTLDHLRFPMEMHIVNIKSTHPNLTSALEDPTGLAVLGVFIDVSSLHNENFHPISSALSFVAYKGQTKPIRPFPLINLLPQNNLAQYYRYHGSLTTPPCSQVVLWTVYEVPIYISWSQFDRFISGIYSTQEDENDQVLLHDNYRHIHPTYSRTVYASKDARRLTNSASCLTLFHLLNVLQLLGLLNL

>Tda|Ca5a

MVTLTAIVSPLGRHIHRHLVRQVRSQRWIPARKCNLSLCSSKLALSQLHPMWQEPLAIPGGDRQSPIDIMVRKSVFDSQLRPLKTQYDPRSCQQIWNNGYSFLVEYDDTNDKSTLKGGPLEDQFRLCQFHFHWGENNAWGSEHSIDRRLYPAELHIVHWNSDKYSLFEEAVVEENGLAVIGVFLKVGKRHEGLQKLVDALPAVRHKDSVVEFNKFDPACLLPKNIDDYWTYAGSLTTPPLTEAVTWIIMKQHIEVSHDQLAVFRSMLFTSAEEQVQRSMVNNFRVQQALKGRSVRSSFTPFLQDAPSME

>Tda|Ca8

MADNVIEESDYYPGKDELEWGYEEGVEWGLLFPEANGEYQSPINLNSREACYDPRLLEVGLNPNYVVCRDCEVINDGHTVRIMLKSKSVVTGGPLPSDHEYELNEVRFHWGRENQRGSEHTVNFKAFPMELHLIHWNATLFNSVEDAMGKKNGILIIALFVQVGKEHLGLKAITDVLQDLQYKGKTKIIPCFNPNTLLPDPLLRDYWVYEGSLTTPPCSEKVTWILYRYPLTISQMQIEEFRRLRSHVKGAELLEGNDGMLGDNFRPTQPLSDRVVRAAFQ

>Tda|Ca2

MSHGWGYAKHNGPHKWCESFKIANGPRQSPIDIIPESASYDSSLKPLTLKYDPSTSLEILNNGHSFQVTFADDEDSSTLTGGPISGIYRLKQFHFHWGDGDEKGSEHTVDGKCYPAELHLVHWNTKYSSFAEAADKPDGLAVVGVFLEIGAENAQLQKILVALDSIQCKGTQNSFANFDPSVLLPSSLDYWTYPGSLTTPPLYESVSWIVCKESISISSAQMETFRSLRFSEEEEEACCMVNNYRPPQPLKGRAIRASFQ

>Ost_evm.model.001551F_arrow_pilon.9|ca15a

MIVLLVIFAALLCPTVHSVDASVEWCYHNPTCNVTTWSKIVPEYCNGSQQSPINIVTASVQGNPNLTSFNLTGFDDNSTFMSIVNSGKSVEINLDDAKIKVQGGDLPGVYNIKQFHLHWGNGSSSPGSEHTVDGKQYPMELHIVSVHSKYNGLSAALAAKDSTALAVLGFFIEGTDEANKTKSWDILTSYLKNISKSGDQTSDIMNQTTLNGLTEGVDRTKYYRYQGSLTTPSCNEAVIWTVFKEPVKVSYEFINRFSTTVLFKDANASVLNTNSFRGVQPLNGRVVTSQIPSAAPSSSTLSVITVLLLSRLFWL

>Ost_evm.model.001294F_arrow_pilon.1|ca15a

MIVLLVIFAALLCPTVHSADASVEWCYHNPTCNVTTWSKIVPEYCNGSQQSPINIVTASVQGNPNLTSFNLTGFDDNSTFMSIVNSGKSVEINLDDAKIKVQGGDLPGVYNIKQFHLHWGNGSSSPGSEHTVDGKQYPMELHIVSVHSKYNGLSAALAAKDSTALAVLGFFIEGTDEANKTKSWDILTSYLKNISKSGDQTSDIMNQTTLNGLTEGVDRTKYYRYQGSLTTPSCNEAVIWTVFKEPVKVSYEFINRFSTTVLFKDANASVLNTNSFRGVQPLNGRVVTSQIPSAAPSSSTLSVITVLLLSRLFWL

>Ost_evm.model.001333F_arrow_pilon.9|ca15c

MKPTHYIPAYRQANQVREMIVLLVTCLAALLCPTVHSADASVEWCYHKPACNFTTWSKIAPKYCNGSQQSPINIVTASVQGNLNLTSFNLTGFDDNSTFMSIVNSGDSVVVNLDDEKIIVQGGALPGVYNTKQFHLHWGNGSSSPGSEHTVDGKQYPMELHIVSVHSKYNGNVSAALAAKDSTALAVLGFFIEATSDVNKTKSWDILTSYLKSIPSSGDQTSDIMNQITLNGLTEGVDRTKYYRYQGSLTTPSCNEAVLWTVFKEPVKVSQNFINRFSTTVLFKDANASVLNTNNFRGVQPLNGRVVTSQITSAAPSSSTLSVITVLLLSSLCWL

>Ost_evm.model.001293F_arrow_pilon.10|ca15c

MIVLLVIFAALLCPTVHSEDASADFTTWYKIVPEYCNGSQQSPINIVTASVQVNPNLTSFNLTGFDDNSTFMSIVNSGESVVVNLNDAKIKVQGGDLPGVYDTKQFHLHWGNGSSSPGSEHTVDGKQYPMELHILSVHSKYNGNVSAALAAKDSTALAVLGFFIEGTDEANKTKSWDILTSYLKNISKSDQKSDIMNQITLNDLTEGVDRTKYYRYQGSRTTPSCNEAVIWTVFKEPVKVSNEFINRFSTTVLFKDANASVLDTTTSEELPWLPDV

>Gpr|ca15a

MMLLMTCVVAVLLSDFSFAKDWCYFGCENSLPNWGKSYPKCNGRKQSPINIDTQKVIKNQELASFELINFSLPHTMKMLKNNGHTVECELKAGAAGVRGGGLKHKYTVLQFHFHWGGKDLMQHPGSEHSLNGHRSPLEMHIVSRRSDLNDSTAAKVQDGFAVMGFFIEGNKKEKEKEKKTSQVWESFTDYLQKIPRKGDKVRIMEPFSMHQLLKGVDLSKYYRYNGSLTTPPCDEAVVWTIFKDPIRISRELVHANSFLIPNNSTTFYKTFMHDHLPSPNCIGSGSGFFIRHIINYTGYNQ

>Gpr|ca15c

MIVLLVTCLAALLCPTVHSADASVEWCYHKPACNFTTWSKIAPKYCNGSQQSPINIVTASVQGNPNLTSFNLTGFDDNSTFMSIVNSGESVVVNLDEAKIKVQGGALPGVYNTKQFHLHWGNGSSSPGSEHTVDGKQYPMELHIVSVHSKYNGDVSAALAAKDSTALAVLGFFIEGTNEANKTKSWDILTSYLKNISKSGDQTSDIMNQITLNGLTEGVDRTKYYRYQGSLTTPSCNEAVIWTVFKEPVKVSQNFINRFSTTVLFKDANASVLNTNSFRGVQPLNGRVVTSQITSAAPSSSTVSVITVLLLSGLC

WL

>Leu|RH50

MKKKSTNLRVRLPVLIFALEVMIVVLYAFFVTYDDQANALLQNNQTRPMDNSLYQNYPFF

ADIQVMIFLGFGCLLAFFRRYGFGGMVFNFLIATFTIQWAILVQGFFQFYYDGKIHLGVL

NLINAEFACAVVLISFGAVLGKTSPVQLLVMALLEIPVFGVTEWAVLKYLKINDAGGSIL

IHIFACYFGLGVTFVLYRPSINEGHPKEKTSYQSDLLSVIGTLFLWVFWPSFNSALTFKG

DDQHRAVLHTFIGLSASTITAFALSSMLSKNGKISMADVQNVTLAGGVTVGASVDMMISP

AAAFILGILGCIACMLGYKYLSPFLARRLRLQDQCGIHNLHGLTGLISSLAGICGILVAT

EETYGPSLYQTFSHRAPPEGDPLLAELRVFIPDLEAGLGRSAQEQALFQVAAVFGTIAVS

AVGGLLTGVVLKLPYLASPSDENCFDDELFFDVPPDYNSVNGPQEAWSCVLRDANKTDVD
